# Sensorineural regulation of skull healing implicates FGF1 signaling in non-healing bone

**DOI:** 10.1101/2025.11.24.690319

**Authors:** Allison L. Horenberg, Carlos U. Villapudua, Mingxin Xu, Eric Z. Zeng, Shaquielle Dias, Tanishk Sinha, Mary Archer, Erica L. Scheller, Arvind P. Pathak, Aaron W. James, Warren L. Grayson

## Abstract

Bone injuries demonstrate rapid peripheral nerve ingrowth followed by nerve pruning as healing ensues. However, in non-healing bone injuries, peripheral innervation remains elevated, the implications of which remain unknown. Therefore, we investigated the neuroskeletal microenvironment in subcritical-and critical-sized calvarial defects using quantitative 3D fluorescence imaging. We identified elevated densities of peripheral nerves and Osterix positive (Osx+) osteoprogenitors in critical-sized defects, while osteogenic differentiation markers were severely diminished. Moreover, Osx+ osteoprogenitors in critical-sized defects exhibited enhanced proximity to peripheral nerves, which in turn was associated with increased osteoprogenitor cell proliferation. Using retrograde tracing in conjunction with single cell RNA-sequencing of sensory nerves from the innervating trigeminal ganglia, genes encoding for nerve-secreted proliferative and anti-differentiation factors were identified. Specifically, FGF-1/FGFR-1 signaling was identified as a significant neuroskeletal interaction with critical-sized defects demonstrating higher FGF-1 expression in fluorescence imaging. Presence of FGF-1 in neurons innervating the calvarial was confirmed, and neural conditioned media depleted for FGF1 showed enhanced induction of osteogenesis when placed on parietal bone cells. Collectively, we identify a sensorineural-skeletal signaling interaction elevated in critical-sized defects that can be leveraged as a potential therapeutic target for enhancing bone healing.

## INTRODUCTION

The skeleton contains a dense network of peripheral nerves that respond to environmental and mechanical stimuli. Recent studies suggest an important role for peripheral sensory nerves in coordinating osteoprogenitor activity during development and following injury.^1,2^ Conversely, in fracture non-unions, there exist higher densities of peripheral nerves within the defect site than in healing samples indicating a potentially negative correlation between innervation and regeneration.^3,4^ Interestingly, both neurotrophins (e.g. nerve growth factor, NGF) and nerve retraction factors (e.g. Sema3a) have been shown to enhance bone formation after injury.^5,6^ These findings suggest that nerves have the potential to play multiple, possibly contradictory roles following bone injuries that has profound implications for therapeutic strategies targeting neural signaling.

Sub critical- and critical-sized calvarial bone defects are defined by their disparate post-injury responses, with critical-sized injuries demonstrating an inability to heal spontaneously. Imaging the dynamic axonal responses to these injuries could provide insight into the neural contributions responsible for these inherent differences. Previous attempts to do this relied on 2D histologic cross-sections or 3D confocal imaging of small regions-of-interest. The limited fields-of-view inherent to these methods make it challenging to establish a comprehensive and quantitative evaluation of overall nerve ingrowth. However, recent methods such as quantitative light sheet microscopy (QLSM) can be used to visualize nerves and their 3D interactions over large skull volumes.^7,8^ Using advanced analysis methods, QLSM allows quantification of the spatial proximity between filamentous structures, such as nerves, and neighboring cells, such as osteoprogenitors. These spatial associations provide hints into signaling factor interactions that may mediate the healing cascade. Elucidating the nerve-osteoprogenitor cross-talk in the skull is further complicated by its unique anatomy (e.g. nerve cell bodies are located in distal ganglia) which makes it challenging to isolate precise nerve-derived signaling factors. Nerve tracing methods using lipophilic dyes such as Luxol Fast Blue have been effective at labeling nerves that innervate a tissue of interest. Although these dye-based techniques can label specific nerve nuclei, they are incompatible with next generation gene sequencing techniques. In these instances, it is preferable to use viral retrograde tracing that allows label integration for subsequent sequencing and computational analysis. Recently, these methods have been performed in long bones, where skeletal nerve profiles were investigated over multiple timepoints after injury.^9^ This technique has not been employed in the calvaria, which demonstrates bone formation via intramembranous ossification and is innervated by nerves from the trigeminal ganglion, and the precise nerve signature following bone injuries has yet to be explored.

In this study, to elucidate neural dynamics and gain insight into their roles in mediating osteoprogenitor responses during bone healing, we compared the axonal sprouting from peripheral nerves into healing and non-healing skull defects. Specifically, we investigated the neuroskeletal microenvironment in subcritical- (1-mm) and critical-sized (4-mm) calvarial defects in adult C57BL6/J mice. Using advanced 3D fluorescent imaging techniques and quantitative analyses, we identified unique spatial distributions and cell-cell interactions between peripheral nerves and osteoprogenitors, as a function of injury type. To investigate signaling factors that mediated these interactions, we incorporated viral retrograde tracing and single cell RNA-sequencing (scRNA-seq) of cells in the trigeminal ganglia as well as in the naïve calvaria and identified key factors that mediate osteoprogenitor proliferation. To validate whether these interactions impact bone healing, we leveraged scRNA-seq datasets, performed additional immunofluorescent staining, and evaluated in vitro osteogenic assays. Overall, these data provide critical insights into the role of calvarial nerves in mediating bone formation and their differential impact in healing and non-healing calvarial defects.

## METHODS

### Materials

All antibodies used in this study can be found in **Supplemental Table 1.**

### Animal Models

All animal experiments were approved by the Johns Hopkins University Institutional Animal Care and Use Committee (Protocol Nos. MO21M146 and MO24M96). Animals were housed and cared for in Johns Hopkins’ Research Animal Resources central housing facilities. C57BL/6J (The Jackson Laboratory, Stock No: 000664) mice used in this study were 10-14 weeks of age and were all male. B6.129(Cg)-Scn10atm2(cre)Jwo/TjpJ (The Jackson Laboratory, Stock No: 036564) and B6.Cg-Gt(ROSA)26Sortm14(CAG-tdTomato)Hze/J (The Jackson Laboratory, Stock No: 007914) were crossed to generate NaV1.8-tdTomato mice. Mice are maintained by sibling mating and used as NaV1.8+;tdTomato+/+. NaV1.8-tdTomato mice were 10-14 weeks of age and were both male and females as indicated.

### Calvarial Defect Surgery

For calvarial defect surgeries, mice were anesthetized via intraperitoneal injection with ketamine (100 mg/kg) and xylazine (10 mg/kg). Following anesthesia, the paw-pitch test was performed to ensure complete sedation. Buprenorphine-ER (1 mg/kg) was administered subcutaneously. To access the calvarial bone, the mouse head was shaved, sterilized with alcohol and betadine, and draped under sterile conditions. Next, mice were cured in a stereotaxic frame to stabilize the skull. A sagittal incision was made along the midline of the calvaria. Skin and connective tissue were removed from the parietal bones to ensure localization of the surgical site. A 1-mm or 4-mm defect was then created using a burr hole or trephine drill bit, respectively. To ensure similar locations of the inner osteogenic front for all defect sizes, both defect edges were positioned at 1 mm from the sagittal suture, with the 4-mm centered within the right parietal bone. Special precaution was taken to avoid disturbing the underlying dura mater. Following defect creation, the skin incision was closed with 6-0 nylon monofilament sutures. Mice were monitored daily for up to seven days for any neurological deficits, infection, pain, or distress. In addition to uninjured controls, mice were euthanized at 1-, 2-, 4-, and 8-weeks post-surgery.

### Calvarial Harvest

Mice were anesthetized with ketamine (100 mg/kg) and xylazine (20 mg/kg). Following anesthesia, mice were injected subcutaneously with 200 U of heparin prior to perfusion. An incision was first made at the xiphoid process and additional incisions were made to gain access to the rib cage and heart. Following disruption of the pleural cavity, heparinized saline (10 U/mL in 1X PBS) was perfused via a 20G blunt needle into the heart via the left ventricle. Prior to perfusion, the right atrium was cut open to allow an open circulation. Perfusion was performed at a rate of 10 mL/min. Following perfusion, the skin was removed around the skull and the calvaria was harvested while maintaining the periosteum and the dura mater. Calvaria were fixed in 4% methanol-free paraformaldehyde overnight at 4°C. Following fixation, calvaria were washed three times for 1 hour with PBS at RT.

### Trigeminal Ganglia Harvest

Following calvaria harvest, the brain was carefully removed, severing the optical nerves but without disturbing the underlying fascia. The pituitary gland was scraped away and the trigeminal ganglia were identified. To remove the trigeminal ganglia, all three peripheral nerve branches and the central nerve branch were first cut from the body of the ganglia. Next, the underlying connective tissue was carefully cut to release the ganglia. Trigeminal ganglia were fixed in 4% methanol-free paraformaldehyde overnight at 4°C. Following fixation, trigeminal ganglia were washed three times for 1 hour with PBS at RT.

### μCT Imaging and Analysis

Following QLSM, calvaria samples were washed with PBS at RT and embedded in 4% (40 mg/mL) agarose for μCT imaging. Samples were embedded in a shortened 15 mL Eppendorf tube that was cut in half and sealed with parafilm. μCT images were acquired on a Skyscan 1275 scanner (Bruker) with the following acquisition parameters: 0.5 mm aluminum filter, 10 µm pixel size, 55 kV, 145 μA, 335 ms exposure time, 0.5 rotation step, 5 averages, and 180° scan. 3D image reconstruction was performed using the NRecon software (v.1.7.0.4, Bruker).

μCT image analysis was performed using Mimics 14 Software (Materialise). Images were imported and cropped to the defect region. Image contrast was adjusted to exclusively visualize the bone. Next, the reconstructed bone region was masked. Either a 1-mm-or 4-mm-radius cylindrical .STL file was loaded into the workspace and perpendicularly aligned with the defect region. Boolean operations were used to calculate the overlap between the cylinder and the bone and represented the volume of new bone formed within the defect.

### Whole-Mount Immunostaining and Optical Clearing

To stain and image calvaria via QLSM, we used a previously published protocol.^8^ Briefly, samples were blocked overnight at 4°C in a blocking buffer containing 10% V/V normal donkey serum and wash buffer (0.11M Tris, 0.151M NaCl, 0.05% V/V Tween-20, 20% V/V dimethylsulfoxide, pH 7.5). Calvaria were blocked for 8 hours at RT to block endogenous biotin. Primary staining was performed with antibodies for 7 days at 4°C in blocking buffer. Next, samples were stained with fluorophore-conjugated secondary antibodies for 7 days at 4°C. NaV1.8-tdTomato samples were antibody stained and amplified with streptavidin labeling. First, endogenous streptavidin binding was blocked (Thermo Fisher Scientific NC9047822) and amplified with a AF555 streptavidin-conjugate. All antibodies were diluted in blocking buffer. Between each staining incubation, samples were washed five times for at least 1 hour in wash buffer. Following staining and immediately prior to QLSM, samples were cleared in a graded series of 2,2-thiodiethanol (TDE in TBS-Tween; 25%, 50%, 75%, 90%, 100% × 2). Tissue clearing was performed for a minimum of 2 hours at RT or overnight at 4°C. Samples were stored in 100% TDE prior to imaging.

### Light-Sheet Imaging, Processing, and Analysis

For lightsheet imaging, we used a LaVision Biotec Ultramicroscope II that was aligned for the refractive index of TDE. Calvaria were mounted to a sample mount and immersed in a glass imaging cuvette with 100% TDE. Samples were imaged using three separate tile acquisitions: a 3 × 1 tile with double-sided illumination; a 3 × 2 tile with left-sided illumination; and a 3 × 2 tile with right sided illumination. Tiles were overlapped by 15% to facilitate stitching. The following hardware and settings were used for all scans: ×2.5 zoom with a ×2 dipping cap (×5 magnification, 1.3 μm x–y pixel size), 5.5 Megapixel sCMOS camera, 20 ms exposure time, 0.154 numerical aperture, and 2.5 μm z step size. Three channels were imaged using 561, 640, and 785 nm lasers and 620/60, 680/30, and 845/55 emission filters, respectively. Laser intensities were set for each antibody and held constant for each scan.

All image processing and analysis was performed with Imaris® 9.10, Ilastik®, and ImageJ software running on a Dell Precision 7820 Tower workstation. The workstation was equipped with a dual Intel Xeon Gold 6240 processor, 384 GB DDR4 SDRAM (26661MHz speed), 512 GB and 1 TB SATA SSDs, NVIDIA Quadro RTX5000 graphics card (16 GB GDDR6 memory), and Windows 10 Pro for Workstations.

To effectively segment the axons and quantify calvarial nerve density, we used a previously published Ilastik®, ImageJ, and Imaris® workflow,^10^ which allows accurate segmentation of nerves in conditions with varying signal-to-noise ratios.

### Imaris-based Image Conversion and Stitching

To process the raw data for Imaris®, we converted LaVision Biotec raw OME-TIFF files to the Imaris® file format (.ims) using the Imaris® File Converter 9.8. Tiles were manually aligned and stitched using the Imaris® Stitcher 9.8, resulting in a final 3D image. A cuboidal volume-of-interest (VOI) was positioned around the defect region. VOI dimensions were consistent between data for the same defect size. VOI were cropped and exported into ImageJ for channel separation.

### Ilastik®-based Image Segmentation

We segmented nerves (TUBB3+, NeuF+, NaV1.8-tdTomato+) and osteoprogenitors (Osx+) in all images using Ilastik®,^11^ a freely downloadable machine-learning based image segmentation tool. Briefly, once color channels were separated in ImageJ, two z-slices from two replicates were selected from each region and each condition for Ilastik® training. All features were selected for Ilastik® training to account for changes in intensity, shape, and texture. Ilastik® was trained on individual z-slices to identify nerve/osteoprogenitor and background signals, and prediction masks exported for each region-of-interest. The remaining z-slices for each condition were segmented using batch segmentation and the corresponding prediction masks. Following Ilastik® segmentation, additional binary segmentation images were generated in ImageJ. Next, z-stacks were imported back into Imaris® for subsequent analysis.

### Imaris®-based 3D Analysis

Nerve channels were segmented using absolute intensity surface segmentation and a 10^4^ μm^3^ volume filter to eliminate subcellular-sized segments. Blood vessels (stained with CD31 or Emcn) were segmented using surface segmentation with a 10 μm radius for background subtraction and a 10^4^ μm^3^ volume filter to eliminate subcellular-sized segments. Segmentation thresholds were optimized for each stain and kept consistent between sample ages. Volume density measurements were calculated from the nerve or blood vessel volumes and normalized by the volume of the region-of-interest. Osx+ osteoprogenitors were segmented using the spots module that employed a premeasured XY diameter of 6 μm and z diameter of 18 μm.

Next, images were down-sampled by a factor of two in each dimension. Down-sampling allowed further processing of the initial segmentation to identify vascular phenotypes and spatial nerve associations. Binary masks for all stains were generated. A second segmentation (referred to as the down-sampled segmentation) was performed on these masks which included the Imaris® “Split Objects” Surfaces function with a 10 μm seeding point diameter. CD31^hi^Emcn^-^ and CD31^hi^Emcn^hi^ vessels were segmented using the CD31 mask and filtered based upon the absence or presence of masked Emcn signal within each object, respectively. CD31^lo^Emcn^hi^ vessels were segmented based upon the Emcn mask and filtered to remove objects that co-localized with the masked CD31 signal. Spatial association statistics were generated for each vessel phenotype and Osx+ osteoprogenitor by exporting the shortest distance metrics from the down-sampled nerve segmentation data.

### Cryosectioning, Immunohistochemistry, and Cross-Section Imaging

Following all whole-mount imaging, calvaria and trigeminal ganglia were embedded in Tissue-Tek O.C.T. Compound (Electron Microscopy Sciences 62550-01) and flash frozen in liquid nitrogen. Calvaria were sectioned along the coronal direction on a cryostat (Leica CM3050 S) at a thickness of 30 μm. Trigeminal ganglia were sectioned laterally at a thickness of 10 μm. Sections were collected throughout the full defect region or trigeminal ganglia. For immunohistochemistry, slides were blocked and permeabilized in 10% normal donkey serum with 0.1% Triton-X in PBS for 1 hour at RT. Next, slides were stained with primary antibodies in 10% normal donkey serum in TBS with 0.05% V/V Tween-20 (TBS-Tw) at RT. After three washes for five minutes each in PBS, samples were stained with fluorophore conjugated secondary antibodies and DAPI for 1 hour in 10% normal donkey serum in TBS-Tw at RT. After three additional washes in PBS, slides were mounted with cover slips in a solution of 50% glycerol. Tiled z-stack images were acquired with the 10× lens on a Zeiss AxioObserver with Apotome microscope.

For hematoxylin and eosin (H&E) staining, slides were rehydrated for 10 minutes in H_2_O and stained with hematoxylin for one minute. Next, slides were washed five times for one minute in H_2_O and then, briefly dipped in acid ethanol. Following five additional one-minute washes with H_2_O, slides were incubated in eosin for five seconds. Next, samples were dehydrated in a graded series of ethanol from 70% to 100% for one minute each. Samples were incubated in two one-minute xylene steps and then mounted with cover slips using Permount (Thermo Fisher Scientific SP15-100). Thus, all subsequent cross-sectional analysis was performed along the same anatomical inner osteogenic front region for both defects.

### EdU Staining

EdU staining was performed using the Click-iT™ EdU Cell Proliferation Kit for Imaging, Alexa Fluor™ 488 dye (Thermo Fisher C10337). EdU resuspended in saline (5 mg/kg, 1.25 mg/mL) was injected 24 hours prior to tissue harvest. Following tissue harvest and cryosectioning as described above, slides were stained following the manufacturer’s protocol. Primary and secondary antibody staining was performed after EdU amplification and slides were cover slipped and imaged.

### Retrograde Injections

To label neurons that innervate the parietal bone, AAV-PHP.S-tdTomato (Addgene 59462-PHP.S, titer ≥ 1×10¹³ vg/mL) was used as previously described in long bones.^9^ To first validate the kinetics and localization of tracing, a mixture of virus (3.5µl) and Fast Blue (1.5µl, Polysciences 17740-1) was injected into the right parietal bone periosteum. Ipsilateral and contralateral trigeminal ganglia were harvested and fixed at day 3, 7, 14, and 21. Tracing of parietal bone neurons was evaluated using whole mount imaging on a Zeiss LSM 710 confocal microscope. To estimate the number of tdTomato+ neurons per ganglia, trigeminal cells were isolated in single-cell suspensions and analyzed by flow cytometry using an Attune NxT Flow Cytometer.

### Trigeminal Ganglia Isolation

To isolate trigeminal ganglia neurons in a single-cell suspension, ganglia were first isolated whole as described above. Digestion protocols we adapted from previously published protocols.^12^ Each tissue was next incubated in a digestion buffer containing Collagenase A (20 mg/mL, Sigma-Aldrich 10103578001), Dispase II (20 mg/mL, Sigma Aldrich D4693), and Papain (25U/mL, Worthington Biochemical LS003126) for four 30-minute periods. Between each incubation, ganglia were triturated with a glass Pasteur pipette. The pipette tip was flamed between each digestion to sequentially reduce the diameter of the pipette and manually break down the tissue. The Pasteur pipette was washed in a 0.5% BSA solution between each sample to avoid cross-contamination and reduce the cells from sticking to the pipette. Immediately after digestion, samples were filtered through a 40 μm cell strainer and centrifuged at 100g for four minutes at 4°C. Samples were resuspended in Neurobasal A media (1% penicillin/streptomycin (Gibco), 1X Glutamax (Gibco), 2% B27 (Thermo Fisher Scientific)) and layered on top of a 9% Optiprep gradient (Sigma Aldrich D1556) in PBS. Samples were next centrifuged at 100g for 10 minutes at 4°C with an acceleration of 4 and a deceleration of 0. Supernatant was carefully removed, and cells were resuspended for counting.

### Calvaria Isolation

To isolate parietal bone cells in a single-cell suspension, calvaria were isolated as described above, following cervical dislocation. The right parietal bone was carefully dissected to remove the nearby suture regions. The remaining bone was minced and incubated in digestion buffer containing Collagenase I (1 mg/mL, Worthington Biochemical LS004196) and Collagenase II (1 mg/mL, Sigma Aldrich C2-22-1G). Six 30-minute serial digestions were performed, where the resulting digestion buffer was collected and replaced with an equal volume of fresh digestion buffer. The collected buffer was centrifuged at 300g for five minutes at 4°C. The cell pellet was resuspended in complete αMEM (10% FBS (Gibco), 1% penicillin/streptomycin (Gibco)) and stored at 4°C until the subsequent digestions were completed. Following six digestions, samples were filtered through a 40 μm cell strainer and centrifuged at 300g for five minutes at 4°C. Supernatant was carefully removed, and cells were resuspended for counting.

### Single-Cell Sequencing and Bioinformatic Analysis

Following single cell isolation and prior to sequencing, three replicates of both trigeminal ganglia and parietal bone cells were pooled by tissue type. Single cell suspensions were sequenced at the University of Maryland Baltimore Genomics Center. Library preparation was performed using the 10x Genomics Chromium controller following the manufacturers protocol. Pooled cell suspensions were loaded onto a Chromium Single-Cell A chip along with reverse transcription (RT) master mix and single-cell 31 gel beads, aiming for 30,000 cells per channel. Libraries were sequenced on an Illumina NovaSeq6000. CellRanger was used to perform sample demultiplexing, barcode processing, and single-cell gene counting (alignment, barcoding, and unique molecular identifier (UMI) count). Downstream analysis was performed using Seurat. To ensure maximum retention of high-quality cells, QC was performed iteratively. Final cells were filtered to have >1750 and <6000 detected genes, less than 25% mitochondrial transcripts, and less than 25000 UMIs. To integrate datasets, count tables were merged, while maintaining origin identifiers for both tissues and datasets. Normalization was then conducted using the Seurat log normalization function. Prior to feature selection for principal component analysis, the three QC metrics (nFeature, nCount, percent.mt) were regressed to minimize their effect on clustering. Finally, batch effect correction was done using Harmony^13^ with the tissue and dataset of origin as the anchors. Ligand receptor pair analysis was done using LIANA.^14^ We then used the Panther database^15^ to classify and identify ligand targets to those that could be secreted.

### In Vitro Experiments

For all in vitro experiments, trigeminal ganglia neurons were isolated in a single cell suspension as described above. To ensure isolation of early stage osteoprogenitors, parietal bone cells were isolated from the full right and left parietal bone tissue including the sagittal and coronal sutures. Subsequent parietal bone digestion steps and single cell isolation were performed as described above.

### FGF-1 siRNA Knockdown in neuronal cultures

Immediately following neuron isolation, electroporation was performed using nucleofection. Following trigeminal ganglia cell counting, all cells were centrifuged at 300g for 5 minutes at 4°C and the pellet was resuspended in electroporation buffer (P3 solution and supplement, Lonza V4XP-3012) with 0.3 nmol of either control siRNA (Santa Cruz Biotechnology sc-37007) or FGF-1 siRNA (Santa Cruz Biotechnology sc-39445). The cell suspension was then transferred to a 2.0 mm electroporation cuvette and electroporated with the Nucleofector X system using the DR-114 protocol. Following nucleofection, cells were immediately diluted with warm Neurobasal-A complete media and incubated at 37°C for 10 min to allow cell recovery.

### Culture of primary neurons and bone cells

After trigeminal ganglia digestion and siRNA knockdown, trigeminal cells were plated on poly-D-lysine (100 μg/mL) and laminin (10 μg/mL) coated 24-well plates in complete Neurobasal-A (Gibco) media including 5% FBS, 1% penicillin/streptomycin (Gibco), 1X Glutamax (Gibco), and 2% B27 (Thermo Fisher Scientific) with anti-mitotics, 20 μM 5-fluoro-2-deoxyuridine and 20 μM uridine (Sigma-Aldrich). Media was changed after two days of initial culture.

Isolated parietal bone cells were seeded into a collagen (0.1 mg/mL) coated T-25 flask at 180K cells/cm^2^ in αMEM with 20% FBS and 1% penicillin/streptomycin (Gibco) for expansion. αMEM media was changed every two days until 70% confluency. Cells were passaged once and used at passage two for subsequent experiments.

### Neuronal Conditioned Media Collection

For neuron conditioned media collection, media was changed from complete Neurobasal-A to αMEM with 1% FBS and 1% penicillin/streptomycin on day 3 of culture. Conditioned media was collected on days four through six and stored at -80°C until further use. Knockdown efficiency was confirmed with a mouse FGF-1 ELISA (Abcam ab223587).

### Osteogenic Differentiation

Parietal bone cells were seeded into collagen coated 48-well plates (5000 cells/cm^2^) in osteogenic media containing 10% FBS, 1% Penicillin/streptomycin with 10 mmol/L β-glycerophosphate, 50 µmol/L of Ascorbic acid and 100 nmol/L of dexamethasone. In select experiments, the media contained recombinant mouse FGF-1 (R&D Systems 4686-FA) with concentrations ranging from 2 ng/mL to 20 ng/mL. In conditioned media experiments, the base αMEM was replaced with neural conditioned media with either control siRNA or FGF-1 siRNA. For all osteogenic culture experiments, media was changed three times per week. DNA (Quant-iT PicoGreen dsDNA Reagent, Thermo Fisher Scientific P11495) and calcium (Calcium (CPC) LiquiColor Test, Thermo Fisher Scientific SB0150250) assays were performed on independent wells for all experiments by following manufacturer’s protocols.

### Statistics

GraphPad Prism was used to perform all statistical tests. Either an unpaired two-tailed t-test or a one- or two-way ANOVA with a post-hoc Tukey HSD test was performed. P-values < 0.05 were considered significant.

## RESULTS

### Critical-sized calvarial defects demonstrate persistent innervation at later post-injury timepoints

At eight weeks after injury, 4-mm defects demonstrated lower bone volume fraction than 1-mm defects within the defect region (Figure 1A-B). Moreover, 4-mm defects exhibited less than 10 percent bone formation, validating them as non-healing defects.^16^ H&E staining revealed fibrous tissue within the defect centers at eight weeks after injury (Supplemental Figure 1). Visualization of beta-3 tubulin-stained (TUBB3+) nerves within the defects at 1 week after injury revealed a dense infiltration of nerves into both defect sizes (Supplemental Figure 2A, Figure 1C). Nearly all the nerves visualized at week 1 were thinner than at later time points (i.e. week 8) and did not span the full defect diameter. The expansion of nerves was not localized to the defect area but appeared to be widespread throughout the parietal and frontal bones including on the contralateral side for both defect sizes. At weeks 2, 4, and 8, the large nerve infiltration seen at week 1 gradually subsided while thicker, longer nerves were found traversing the full defect region.

**Figure 1:**
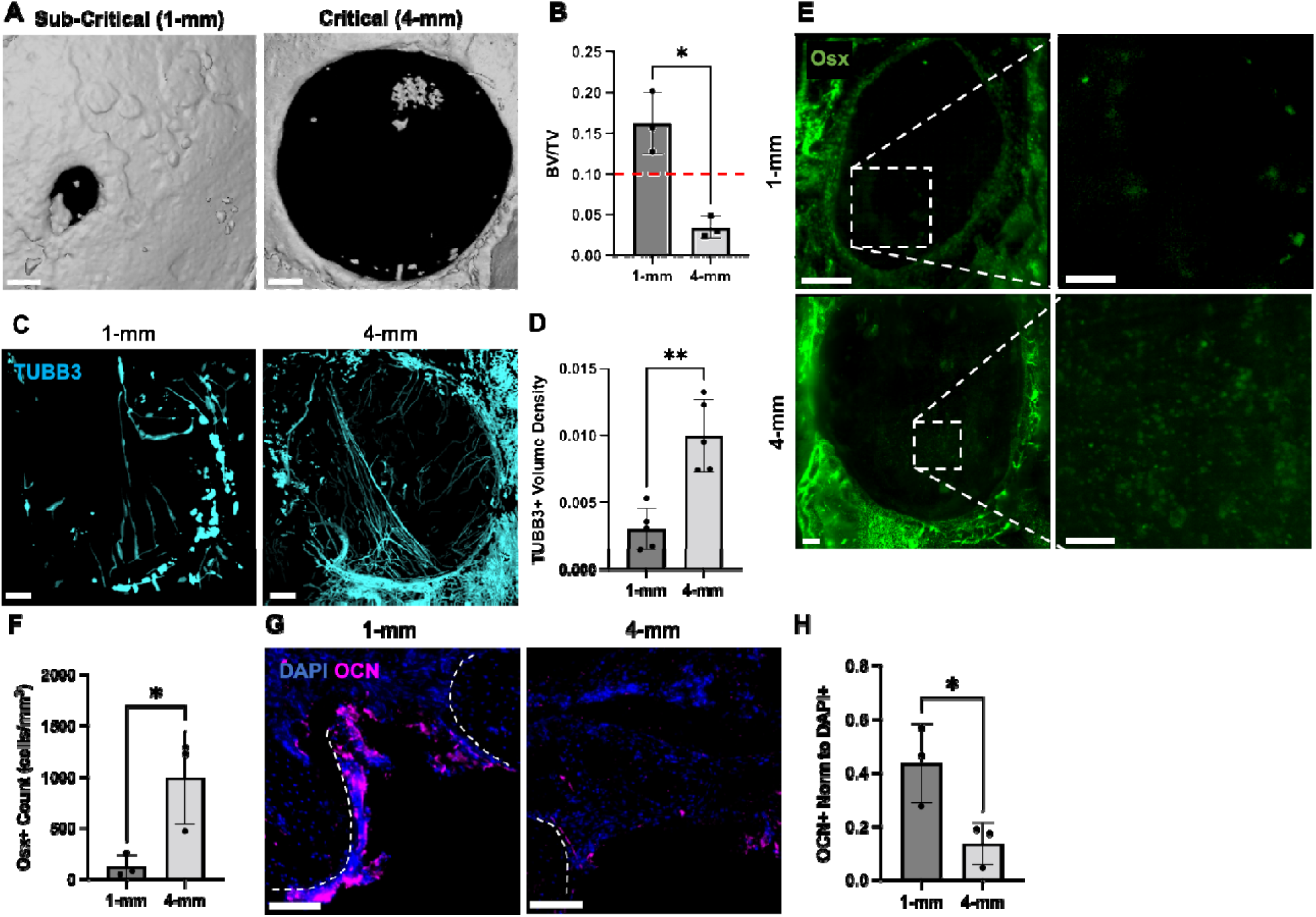
Critical-sized defects demonstrate hyper-innervation and elevated osteoprogenitor density at 8 weeks following injury. **A**) 3D μCT images for sub-critical (1-mm) and critical (4-mm) calvarial defects. Scale bar: 500 μm. **B**) Bone volume (BV) quantification normalized by total defect volume (TV). Dash red line demonstrates 10% healing. **C**) Maximum intensity projection (MIP) images of TUBB3+ nerves acquired with QLSM. 1-mm scale bar: 200 μm. 4-mm scale bar: 300 μm. **D**) Beta III Tubulin (TUBB3+) volume density quantification. **E**) MIP images of Osterix (Osx+) osteoprogenitors acquired with QLSM. Dashed lines represent zoomed regions on the right. 1-mm scale bar: 200 μm. 4-mm scale bar: 300 μm. Zoomed region scale bar: 100 μm. **F**) Osx+ osteoprogenitor count quantification. **G**) MIP images of Osteocalcin (OCN – magenta) and DAPI (blue) from 30 μm calvarial cross-sections from sub-critical (1-mm) and critical (4-mm) calvarial defects. Region-of-interest includes the medial osteogenic front of the defect. Dashed lines represent bone edge. Scale bar: 100 μm. **I**) OCN+ area normalized to DAPI area quantification. Data are Mean ± 1 SD. p* < 0.05. p** < 0.01.

At week 8, 4-mm defects contained a larger volume density of TUBB3+ nerves than 1-mm defects (Figure 1D; Supplemental Figure 2B). In both the 3D images and histologic cross-sections, we observed large nerve sections in the 4-mm defects, which were not found in 1-mm defects. Similar to TUBB3+ imaging, we found a higher density of sensory Neurofilament-stained (NeuF+) nerves in 4-mm defects at 8 weeks post-injury as compared to 1-mm defects (Supplemental Figure 3A-B) and further confirmed these findings in NaV1.8-tdTomato reporter mice (Supplemental Figure 3C-E).

### Type H vasculature and osteoprogenitor cells are elevated in critical-sized defects

Given the known associations of nerves with blood vessels and osteoprogenitors, we next investigated the densities of other cell types in the defect region. As observed previously,^8,17^ CD31^hi^Emcn^hi^ blood vessels constituted the primary vascular phenotype that infiltrated calvarial defects (Supplemental Figure 4A-B). While CD31^hi^Emcn^hi^ blood vessel volume density was significantly elevated in critical-sized defects, there was no statistically significant difference in the spatial associations between nerves and blood vessels in 1-mm and 4-mm defects (Supplemental Figure 4C). Osteoprogenitors identified by immunohistochemical staining for Osterix (Osx) were also found to be elevated in 4-mm defects at eight weeks following injury (Figure 1E-F). Interestingly, a large population of Osx^lo^ cells were found throughout the center of the 4-mm defect region, while limited Osx^lo^ cells were identified in the 1-mm defect. Given this large population of early osteoprogenitors with limited bone formation, we next evaluated the expression of bone differentiation markers. Osteocalcin (OCN) is a marker of osteoblast differentiation and is secreted into the bone matrix. We found a significantly lower OCN area in 4-mm defects (Figure 1G-H) indicating that the higher density of osteoprogenitors did not necessarily result in enhanced expression of differentiation markers.

### Osteoprogenitors exhibit unique spatial interactions with nerves in critical-sized calvarial defects

Due to the varied intensities of Osx+ expression in osteoprogenitors between the edge and the center of the defect region, we used an Ilastik® pixel classification workflow to better segment these cells. We used NaV1.8-tdTomato mice to investigate the neuroskeletal interactions between nerves and Osx+ cells throughout the center of 1-mm and 4-mm defects (Figure 2A). We identified a distinct spatial association between nerves and Osx+ osteoprogenitors in 4-mm defects with a large fraction of them residing within 10 μm of nerves (Figure 2B). This fraction of nerve-associated Osx+ cells was significantly higher in the 4-mm defects compared to the 1-mm injuries (Figure 2C) and interestingly, corresponded with increased clustering (i.e. spatial association) of osteoprogenitors to their three nearest neighbors (i.e. adjacent osteoprogenitors) (Figure 2D). When Osx+ osteoprogenitors were stratified based on their spatial association with nerves (<10 μm distance), we identified uniquely enhanced clustering in nerve-associated osteoprogenitors in 4-mm defects (Figure 2E). This trend was confirmed for up to nine nearest neighbors (Figure 2F). Notably, when stratifying between male and female mice, we did not identify any differences between the nerve and osteoprogenitor density, neuroskeletal association, or osteoprogenitor clustering (Supplemental Figure 5). Thus, higher cellular density of Osx+ osteoprogenitor cells was observed in association with sensory nerve axons in the context of non-healing skull defects.

**Figure 2:**
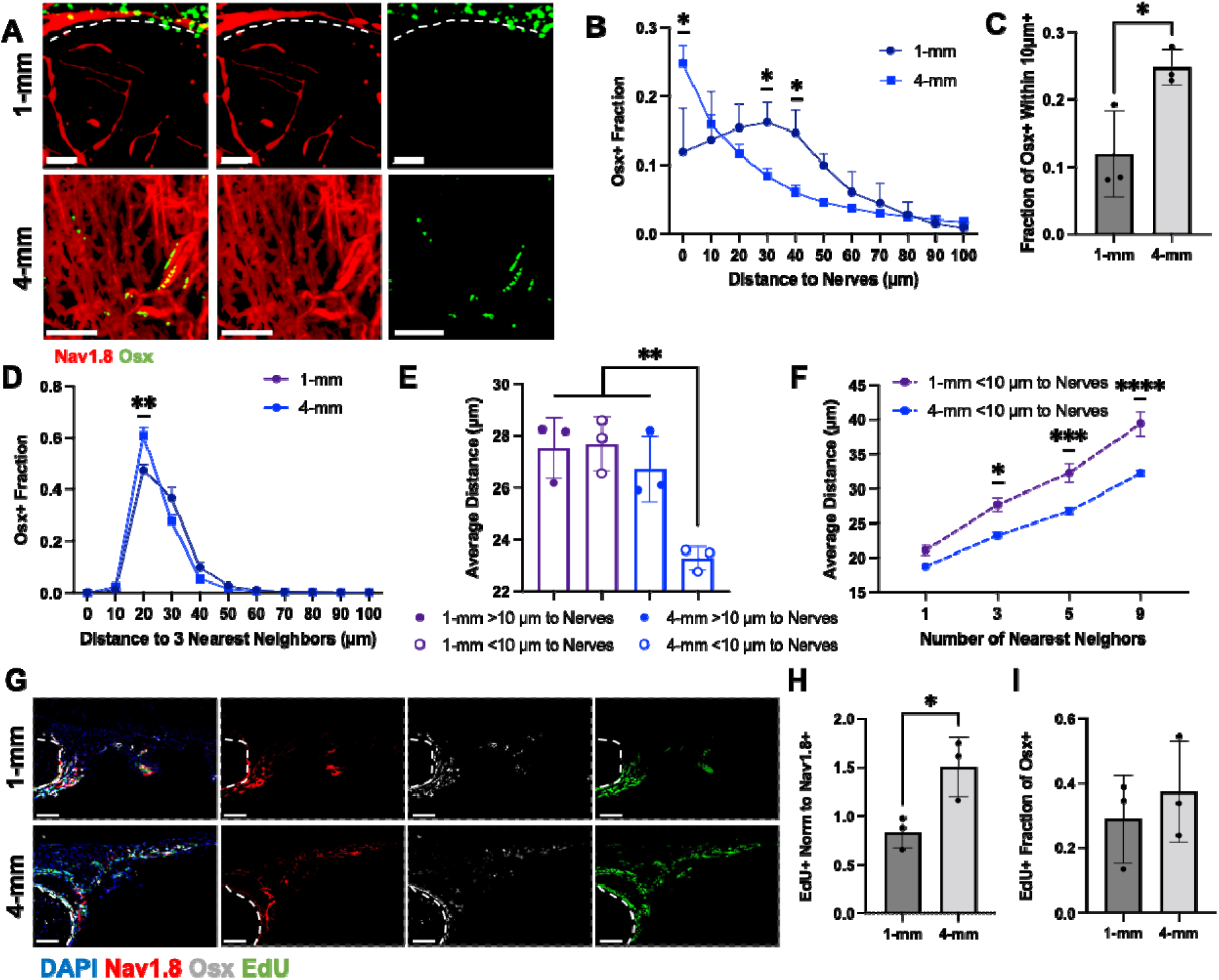
Osteoprogenitors are spatially associated with nerves within critical-sized defects. **A**) MIP image of NaV1.8-tdTomato+ nerves (red) and Osx+ osteoprogenitors (green) within 1-mm and 4-mm defect regions acquired with QLSM. The dashed line represents the bone edge. Scale bar: 200 μm. **B**) Histogram of the spatial association of Osx+ osteoprogenitors and NaV1.8-tdTomato+ nerves. **C**) Quantification of the fraction of Osx+ osteoprogenitors within 10 μm of nerves. **D**) Histogram of Osx+ osteoprogenitor distance to its three nearest neighbors. **E**) Quantification of the average Osx+ osteoprogenitor distance to its three nearest neighbors for subsets at >10 μm and <10 μm from nerves reveal the unique association in 4-mm injuries. **F**) Quantification of the average Osx+ osteoprogenitor distance to one, three, five, and nine nearest neighbors for subsets at <10 μm from nerves. **G**) MIP image of NaV1.8-tdTomato (red), Osx+ osteoprogenitors (white), EdU (green), and DAPI (blue) from 30 μm calvarial cross-sections from sub-critical (1-mm) and critical (4-mm) calvarial defects. Region-of-interest includes the medial osteogenic front of the defect. Dashed lines represent the bone edge. Scale bar: 100 μm H) EdU+ area normalized to NaV1.8-tdTomato+ area quantification. I) EdU+ area fraction of Osx+ area quantification. Data are Mean ± SD. p* < 0.05. p** < 0.01. p*** < 0.001. p**** < 0.0001.

One potential mechanism underlying the higher Osx+ cell density would be sensory nerve-derived mitogens. In order to investigate this possibility, EdU labeling was performed in NaV1.8 reporter mice 24 hours prior to harvest. Following calvaria cross-sectioning, NaV1.8-tdTomato samples were stained for EdU and Osx (Figure 2G). When normalized to the NaV1.8-tdTomato nerve density contained within the cross section, significantly more EdU+ proliferative cells were found in 4-mm defects then 1-mm defects on a per nerve basis (Figure 2H). When only Osx+ cells were considered, there was no significant difference in proliferation between defects (Figure 2I).

### Calvarial nerves express high levels of FGF-1 for osteoprogenitor signaling

Prior to investigating the nerve to osteoprogenitor signaling, we validated the kinetics and specificity of the AAV-PHP.S-tdTomato and Fast Blue retrograde tracing from the right parietal bone to the ipsilateral trigeminal ganglia (Supplemental Figure 6A). The tdTomato+ signal intensity in the trigeminal ganglia stabilized by day 14 and remained constant at day 21 (Supplemental Figure 6B). Following scRNA-seq of parietal bone and innervating trigeminal ganglia, followed by quality control, and Harmony integration, we identified a total of 33,449 cells, including 19,559 parietal bone and 13,890 trigeminal ganglia cells (Figure 3A). After normalization and dimensionality reduction, we identified 17 primary clusters including various immune populations such as Monocytes/Macrophages (Msr1, Adgre1), Granulocytes (Cxcr2), Neutrophils (Fcnb), T Cells (Cd3e), B Cells (Pax5), Erythrocytes (Hbb-bt). Additionally, we found various non-immune populations including mesenchymal cells (Col1a1, Pdgfra), endothelial cells (Pecam1, Cdh5), Schwann cells (Gjb1, Cadm2), and satellite glial cells (Aqp4), with mesenchymal cells demonstrating similar fractions to previous studies in naïve bone.^18^ Lastly, we found a small cluster of neuronal cells marked by the expression of (Trpv1, Scn11a, and Mrgprd) (Supplemental Figure 7A). Next, we evaluated the degree of tdTomato+ neuronal labeling and observed that 3.9% of the collected neurons labeled with tdTomato (Figure 3B). Comparing the differentially expressed genes between tdTomato+ and tdTomato- neurons with a specific focus on secreted and membranous ligands, we identified 46 differentially expressed genes, including Notch4, Nrtn, Edn1, Wnt10a, Pthlh, and Lif, which have been positively associated with bone formation,^19–24^ and Cxcl9, which has been negatively associated with bone formation (Figure 3C).^25^

**Figure 3:**
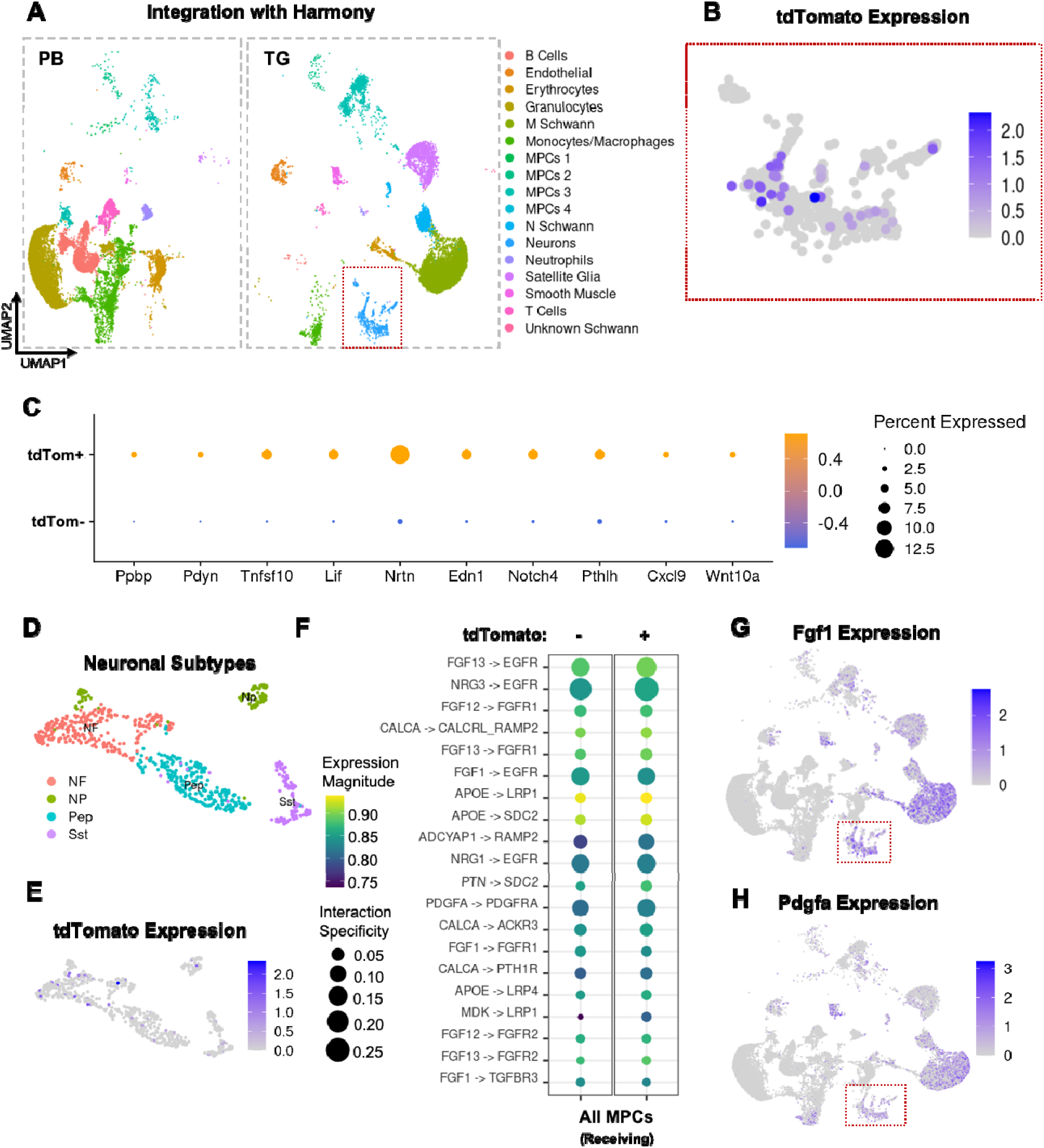
Calvarial nerves express ligands from the FGF, PDGF, and neuropeptide families. **A**) UMAP split by cells from origin of collection. PB = Parietal Bone; TG = Trigeminal Ganglia. **B**) FeaturePlot identifying neuronal cells innervating the parietal bone region via positive expression of tdTomato. **C**) The top 10 differentially expressed ligands between tdTomato+ and tdTomato- neurons. **D**) UMAP illustrating the different neuronal subpopulations. **E**) FeaturePlot depicting enrichment of tdTomato expression within these neuronal subpopulations. **F**) LIANA ligand- receptor pair analysis between tdTomato+ and tdTomato- neurons to combined MPCs. **G**) FeaturePlot depicting Fgf1 expression. Red dashed lines represent neuronal populations. **H**) FeaturePlot depicting Pdgfa expression. Red dashed lines represent neuronal populations.

To further stratify the neuron population, we identified four sub-clusters of neurons including peptidergic neurons, neurofilament (NF)+ neurons, non-peptidergic neurons, and Sst+ nociceptors (Figure 3D, Supplemental Figure 7B). Consistent with our prior observations, we found that most of the tdTomato+ labeling was restricted to NF+ and peptidergic neurons (Figure 3E).^9,26,27^

To evaluate nerve signaling that may mediate osteoprogenitor proliferation, we performed ligand-receptor analysis from tdTomato+ and tdTomato- nerves to all mesenchymal progenitor cell (MPC) clusters using LIANA (Figure 3F). Of the top 20 interactions, there were 11 unique ligands expressed by tdTomato+ neurons and 13 unique receptors expressed by MPCs. FGF12 and 13 were not included for further analysis, as their functions are intracellular. Of the ligands, CALCA, FGF1, APOE, PTN, PDGFA, and MDK have known roles in mediating bone formation.^28–33^ Only CALCA, PDGFA, and FGF1 have been explored in the context of skeletal cell proliferation.^34–36^ However, only PDGFA and FGF1 have been implicated in potentially inhibiting osteogenic differentiation.^36,37^ FeaturePlots for FGF1 and PDGFA (Figure 3G-H) indicated that in the neuron population, FGF1 was more highly expressed than PDGFA in most of the cells. Moreover, within the neuron subtypes, FGF1 was more highly expressed in peptidergic and NF+ neurons (Supplemental Figure 7C-D). When considering ligand-receptor interactions for neuron subtypes, FGF1 and its cognate receptor, FGFR1, were ranked highly in peptidergic and NF+ neurons, but not in non-peptidergic and Sst+ neurons, while PDGFA was ranked highly among all subtypes (Supplemental Figure 7). Therefore, we selected FGF1-FGFR1 signaling for subsequent evaluation due to its higher expression and enhanced specificity for skeletal-innervating sensory neurons.

### FGF-1 expression is elevated in critical-sized defects

Differences in FGF1 expression were assessed in trigeminal ganglia harvested from 1-mm and 4-mm calvarial defects at 8 weeks post-injury using immunostaining (Figure 4A). Although all trigeminal ganglia showed neuronal staining for FGF1, we found significantly higher FGF1 immunoreactivity in trigeminal ganglia from 4-mm defects, when normalized to either NaV1.8-tdTomato+ or TUBB3+ neurons (Figure 4B-C), suggesting that not only are more sensory nerves present within critical-size 4-mm defects, but that there may also be higher sensorineural expression of FGF1. We integrated two previous datasets with our uninjured parietal bone data^38,39^ to evaluate the kinetics of FGFR1 expression after injury. We incorporated an additional 24,336 cells into the full dataset (week 1 = 12,421 cells, week 2 = 11,915 cells) (Figure 4D). Fgfr1 expression was highest in the mesenchymal progenitor cell (MPC) population, while only smooth muscle or neutrophil populations exhibited any additional levels of expression (Figure 4E). Fgfr2 was expressed primarily in the neutrophil population, while Fgfr3 and Fgfr4 were not highly expressed in any cluster (Supplemental Figure 8). Dot plots for Fgfr1, Fgfr2, and Pdgfra stratified based on harvest timepoint revealed that only Fgfr1 expression peaked immediately after injury (Figure 4F) coinciding with the highest density of nerves in 1-mm defects (Supplemental Figure 2B). Given the fact that all Osx-expressing cells within the defect region were Osx^lo^, we stratified the MPC population based on Osx expression (Supplemental Figure 9A-F). This population was further stratified by Runx2 expression, as Osx^lo^Runx2^hi^ cells were predominantly found after sub-critical defect injury by Bixel et al.^38^ Using these stratifications, Osx^lo^Runx2^hi^ cells were one of the primary populations expressing Fgfr1 during all timepoints evaluated (Supplemental Figure 9G). We next investigated the level of FGFR1 expression between 1-mm and 4-mm defects to evaluate changes in expression between defect sizes. Stained NaV1.8-tdTomato 1-mm and 4-mm defects harvested eight weeks after injury for FGFR1 non-significant trend toward increased FGFR1 in 4-mm defects when normalized to NaV1.8-tdTomato density (Figure 4J).

**Figure 4:**
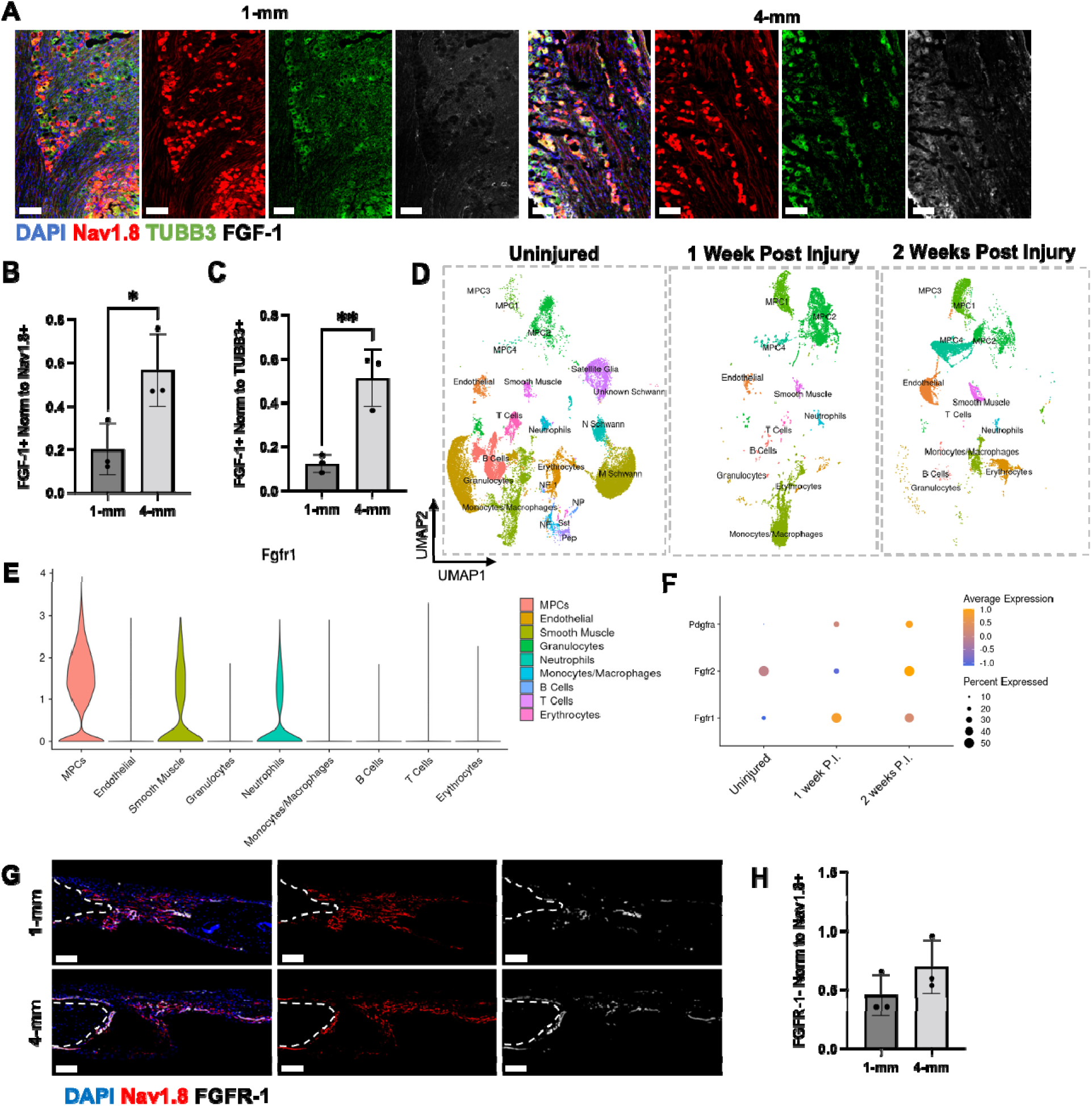
Calvarial defects demonstrate enhanced FGF-1/FGFR-1 signaling. **A**) Trigeminal ganglia (10 μm) cross sections with NaV1.8--tdTomato (red), TUBB3 (green), FGF-1 (white), and DAPI (blue) staining for sub-critical (1-mm) and critical (4-mm) calvarial defects. **B**) FGF-1 area normalized to NaV1.8-tdTomato+ area. **C**) FGF-1 area normalized to TUBB3+ area. **D**) UMAPs of scRNA-seq data from integrated calvaria split by timepoint (week 1 and week 2) following sub-critical sized bone injury. **E**) FGFR-1 expression in each cluster across all integrated timepoints. **F**) Receptor expression of the top ligands identified across timepoints. **G**) MIP image of NaV1.8-tdTomato (red), FGFR-1+ cells (white), and DAPI (blue) from 30 μm calvarial cross-sections from sub-critical (1-mm) and critical (4-mm) calvarial defects. Region-of-interest includes the inner osteogenic front of the defect. Dashed lines represent bone edge. Scale bar is 100 μm H) FGFR-1+ area normalized to NaV1.8-tdTomato+ area quantification. Data are Mean ± SD. p* < 0.05. p** < 0.01.

### Nerve-derived FGF-1 enhances osteoprogenitor proliferation but inhibits bone mesenchymal cell differentiation

Next, we utilized in vitro culture to assess the impact of nerve-derived FGF1 signaling on bone mesenchymal cells isolated from the parietal bone. Parietal bone mesenchymal cells cultured in osteogenic media exhibited no difference in DNA content between 7 and 14 days suggesting that parietal bone cells primarily proliferate in the early days of culture (Figure 5A). However, there was a significant increase in calcium content from day 7 to 14, indicating that calcium deposition primarily occurred after day 7 (Figure 5B). Thus, we hypothesized that the impact of FGF1 on bone cell proliferation and differentiation can be uniquely studied by evaluating early (<D7) and late (>D7) timepoints of culture. Isolated parietal bone cells treated with recombinant mouse FGF1 (rmFGF1) (2 ng/mL – 20 ng/mL) in osteogenic media for 14 days exhibited a dose-dependent increase in DNA content, signifying an increase in bone cell proliferation, with 20 ng/mL rmFGF1 treatment exhibiting the highest DNA content (Figure 5C). Thus, high concentrations of FGF1 increased parietal bone cell proliferation. We next performed Fgf1 siRNA knockdown in isolated trigeminal neurons and collected conditioned media from days 4 - 6 of in vitro neuron culture (Figure 5D). We confirmed Fgf1 knockdown in conditioned media as compared to control siRNA via ELISA (Figure 5E). To investigate whether late knockdown of nerve-derived FGF1, enhances osteogenic differentiation by releasing differentiating cells from their stem-like proliferative state, we cultured isolated parietal bone cells in osteogenic conditioned media with either control or Fgf1 siRNA conditioned media. Treatment began on either day 0 or day 7. We saw no differences in proliferation between control or Fgf1 siRNA samples with conditioned media treatment at either day 0 or day 7 (Figure 5F). However, we identified a significant increase in calcium content in Fgf1 siRNA samples with conditioned media addition at both day 0 and day 7 (Figure 5G). Thus, these data implicate sensory nerve-derived FGF-1 as positively regulating osteoprogenitor cell proliferation while inhibition osteodifferentiation, implicating it as a molecule of potential importance in non-healing bone injuries.

**Figure 5:**
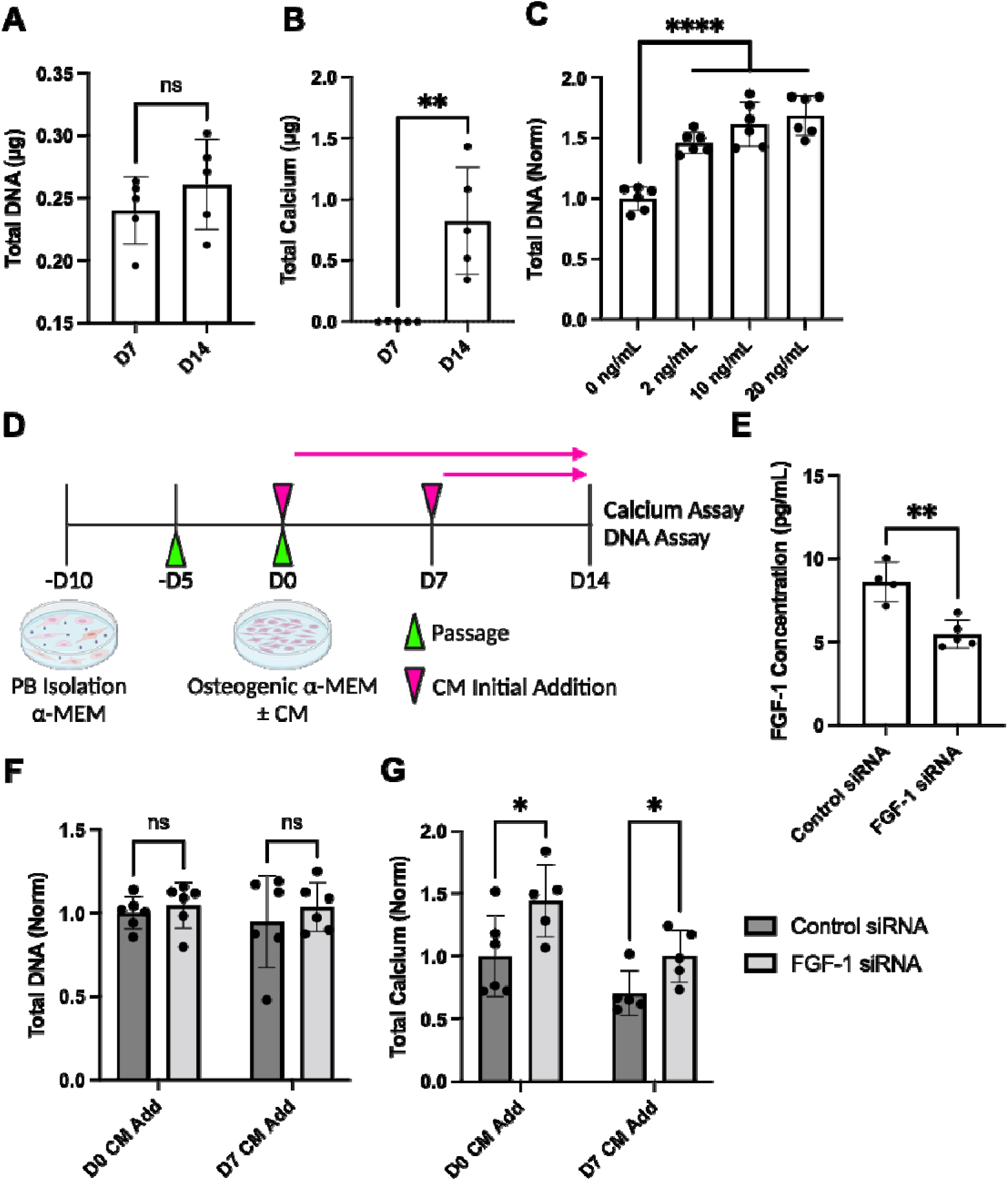
FGF-1 enhances proliferation but inhibits differentiation of parietal bone mesenchymal cells. **A**) Total DNA quantification of parietal bone (PB) cells at day 7 and day 14 of osteogenic culture. **B**) Total calcium quantification of PB cells at day 7 and day 14 of osteogenic culture. **C**) Total DNA quantification of PB cells at day 14 of osteogenic culture with titrated FGF-1. **D**) Schematic of neuronal conditioned media (CM) osteogenic culture. **E**) FGF-1 concentration quantification in CM after neuron transfection with control siRNA or *Fgf1* siRNA. **F**) Total DNA quantification of PB cells at day 14 of osteogenic culture with neuronal CM derived from neurons transfected with control siRNA or *Fgf1* siRNA added at either day 0 or day 7 of osteogenic culture. **G**) Total calcium quantification of PB cells at day 14 of osteogenic culture with neuronal CM derived from neurons transfected with control siRNA or *Fgf1* siRNA added at either day 0 or day 7 of osteogenic culture. Data from C, F, and G are normalized to the control and derived from two independent experiments. Data are Mean ± SD. p* < 0.05. p** < 0.01. p*** < 0.001. p**** < 0.0001.

## Discussion

In this study, we sought to investigate the fundamental differences in the neural responses to bone healing in sub-critical and critical-sized injuries. Using a combination of tissue clearing, quantitative 3D imaging, and novel multi-tissue transcriptomics, we probed whether neural signaling might be an effective therapeutic target for promoting bone healing in critical-sized craniofacial injuries. Immediately following calvarial bone injuries, there was significant axonal sprouting into the defect area followed by gradual retraction or pruning over the next several weeks. This aligns with prior observations in long bones.^1^ However, in critical-sized, non-healing calvarial defects, the densities of peripheral nerves did not return to baseline levels as they did in subcritical-sized healing defects. Interestingly, sustained elevated nerve densities have been correlated with fracture non-unions in long bone. ^3,4^ Nevertheless, a causal relationship has not yet been established, i.e. it remains unclear whether nerves persist because bone has not healed or vice-versa. Based on previous studies, we speculate that environmental factors associated with non-healing fractures, such as mechanical cues, neurotrophic factors, and/or a lack of osteoblast-mediated axonal retraction signals may be responsible for nerve persistence. Firstly, non-healing defects are known to inherently have a softer matrix, with reduced mechanical integrity due to reduced secretion of bone matrix factors into the defect region.^40^ Additionally, nerves demonstrate enhanced growth patterns on softer matrices, resulting in longer axon growth and enhanced neural secretions.^41^ Thus, stiffening of the bone matrix in the sub-critical injuries, may induce gradual nerve retraction, and its lack in critical-sized injuries may result in nerve persistence. Alternatively, non-healing injuries demonstrate elevated inflammatory states, resulting in high levels of macrophage infiltration.^42^ Macrophage-derived NGF is a key factor in mediating neural ingrowth into defects, and persistent inflammation in non-healing bone defects can stimulate nerve persistence. Lastly, while neurotrophins have been shown to encourage nerve axon growth,^6^ other factors acting in tandem are responsible for spatially patterning bone by causing axon retraction.^43,44^ Such axon retraction factors include Sema3a, secreted by osteoblasts.^43,44^ Thus, the lack of bone cell differentiation in critical-sized defects, results in low levels of osteoblasts and limited secretion of Sema3a. We expect that future studies will elucidate the mechanistic underpinnings of nerve persistence in non-healing injuries and help identify novel therapeutic targets.

In addition to elevated nerve densities in critical-sized defects, we also identified elevated levels of Type H vasculature and Osx+ osteoprogenitors. Both vasculature and osteoprogenitors have been closely associated with peripheral nerves. During development, peripheral nerves and blood vessels respond to similar growth signals, resulting in coordinated development and close spatial associations.^45–47^ Bi-directional communication mediates this interaction, with nerves coordinating vessel maturation via secretion of VEGF and established blood vessels coordinating peripheral nerve alignment.^45,47,48^ In bone, nerves are often aligned with blood vessels, and found most frequently within 10 μm of Type H vessels in the uninjured calvaria.^7,49^ Inhibition of TrkA, the receptor for nerve growth factor, demonstrates subsequent reduction of vasculature in bone development and healing, which may be due to the loss of neurovascular signaling.^1,2^ While we did not identify any unique neurovascular signaling interactions in this study, it remains important for tissue homeostasis and response to injury. Therefore, further investigation into neurovascular signaling factors that may play roles in these processes is warranted. As for osteoprogenitors, we visualized an elevated population of Osx^lo^ osteoprogenitors within critical-sized defects. We hypothesize that these cells are early osteoprogenitors, given the fact that similar Osx^lo^Runx2^hi^ cells were identified within the calvarial defect region in a previous study.^38^ As for neuroskeletal signaling to osteoprogenitors, multiple studies have identified a close spatial relationship between nerve patterning and osteoprogenitor location.^50,51^ Regardless of close spatial patterning that may occur between nerves and Type H vasculature or osteoprogenitors, additional mechanisms can mediate the rise in these cell types in critical-sized defects. Similar to nerves, vascular density peaks shortly after injury, which is followed by vascular pruning that occurs along with healing.^52^ Given the lack of healing in critical-sized defects, vascular pruning may be inhibited by nerve-independent mechanisms. In addition, recent studies have illustrated the potential for Schwann cell differentiation into osteoprogenitors.^53^ Thus, elevated nerve persistence, which is accompanied by an increase in Schwann cells, could result in enhanced Schwann cell-derived osteoprogenitors or osteoblasts, although this has not been seen in our recent studies in long bone.^9^

QLSM imaging facilitated the observation of unique osteoprogenitor clustering and enhanced proliferation in spatial proximity to nerves in critical-sized defects. Given the low Osx+ expression of the cells within the defect region, we hypothesized that nerve-mediated proliferation may be occurring in more naïve osteoprogenitors, which may account for the general increase in proliferation seen in 4-mm defects. Previous studies have suggested that nerves mediate proliferation in several contexts including during tooth homeostasis and development, calvarial suture formation, and long bone fracture.^9,50,54,55^ For example, TrkA inhibition resulted in premature sagittal suture thinning, due to dysregulation of BMP and FGF pathways and a loss of nerve-derived FSTL-1 signaling.^55^ This dysregulation reduced progenitor cell stemness which was maintained along the midline of the suture, causing increased differentiation and leading to excess bone formation along the osteogenic fronts. Therefore, to provide some mechanistic insight into the nerve-osteoprogenitor crosstalk, we performed retrograde tracing and scRNA-seq of the parietal bone and the ipsilateral trigeminal ganglia. While both of these tissues have been widely studied independently with scRNA-seq,^56,57^ prior studies have not investigated the signaling factors between the two tissues. Traditionally, studying neural signaling is challenging due to the location of nerve cell bodies in distant ganglia. However, viral retrograde tracing overcomes these limitations by labeling nerve cell bodies innervating the tissue of interest, allowing isolation of tissue-specific nerve signaling factors from scRNA-seq data. These methods were previously used in a long bone fracture model, which identified the key signaling interactions, such as FGF-9 signaling, which mediated^9^ osteoprogenitor proliferation after fracture injury. Of note, both Fstl1 and Fgf9 are expressed in our dataset within trigeminal ganglia neurons. While not directly assessed here, it is possible that FGF1 in combination with these other previously identified mitogenic factors may play a combinatorial or synergistic role in skeletal cell proliferation in a context dependent manner.

Using multi-tissue transcriptomics, we identified highly ranked interactions between skeletal nerves and calvarial mesenchymal cells. In particular, we identified FGF-1/FGFR-1 signaling as a key interaction guiding neuroskeletal signaling and validated enhanced FGF-1/FGFR-1 signaling in critical sized defects. Furthermore, we demonstrated that FGF-1 enhanced osteoprogenitor proliferation, while nerve-derived FGF-1 inhibition at late timepoints of osteogenic differentiation enhanced calcium deposition. Thus, nerve-derived FGF-1 may function by encouraging osteoprogenitor proliferation but inhibiting osteogenic differentiation in critical-sized defects. It is well known that fibroblast growth factors mediate proliferation^58^ and, in bone, function to maintain homeostasis and response to injury.^29^ Interestingly, after nerve injury alone, nerve-derived FGF-1 secretion was also upregulated.^59^ Additionally, in a study of tooth mesenchymal stem cell homeostasis, nerve-derived FGF-1 signaling was shown to mediate Gli1+ progenitor proliferation by controlling the mTOR/autophagy axis.^50^ However, when human bone marrow stem cells (hBMSCs) were treated with FGF-1 in vitro, decreased osteogenic differentiation was observed, seemingly by activating the ERK1/2 pathway.^37^ Also, when hBMSCs were treated with an ERK1/2 inhibitor at the beginning of osteogenic differentiation, both proliferation and calcium deposition were reduced, while treatment during late timepoints of osteogenic differentiation, enhanced calcium deposition with no impact on proliferation.^60^ These results mirror and further validate our findings regarding nerve-derived FGF-1.

Taken together, we have demonstrated a unique role for peripheral nerves in critical-sized calvarial defects. We have combined advanced 3D imaging and transcriptomic analysis to elucidate key signaling factors controlling osteoprogenitor proliferation and differentiation. These factors have the potential to serve as therapeutic targets that can be temporally activated and inhibited throughout the bone healing process. Finally, our scRNA-seq data contains a wealth of information including other factors that may have similar impact on osteogenic differentiation. Future studies can investigate these targets, such as PDGFA and CGRP, in conjunction with FGF1 to enhance bone formation after injury and improve healing outcomes for patients with critical-sized craniofacial injuries.

## Acknowledgements

This work was supported by funding from NIH/NIDCR (1R01DE027957 (WLG), F31DE033910 (ALH), R21DE032420 (ELS), R01DE031488 (AWJ), R01DE031028 (AWJ)), NIH/NIAMS (R01AR079171 (AWJ), R21AR078919 (AWJ)), NIH/NIDDK (R01DK132073), Maryland Stem Cell Research Fund (2022-MSCRFV-5782), the NSF GRFP, NCI (5R01CA237597-05, 2R01CA196701-06A1), and the Orthoregeneration Network. Lightsheet imaging was performed at JHU’s Integrated Imaging Center. Single cell RNA-sequencing was performed at the University of Maryland Genomics Center.

## Author Contributions

A.L.H, A.W.J., and W.L.G. conceived the study. A.L.H., C.U.V., E.Z.Z., S.D., and T.S. performed experiments and analysis. C.U.V. performed computational analysis. M.A. performed mouse breeding and genotyping. E.L.S., A.P.P., A.W.J., and W.L.G. provided experimental insight and direction. A.L.H. wrote the manuscript. All authors reviewed the manuscript and discussed the work.

