## Supplemental Figures for "Sensorineural regulation of skull healing implicates FGF1 signaling in non-healing bone"

400 N. Broadway

Smith 5023

Baltimore, MD, 21231

**Supplemental Figures and Tables**

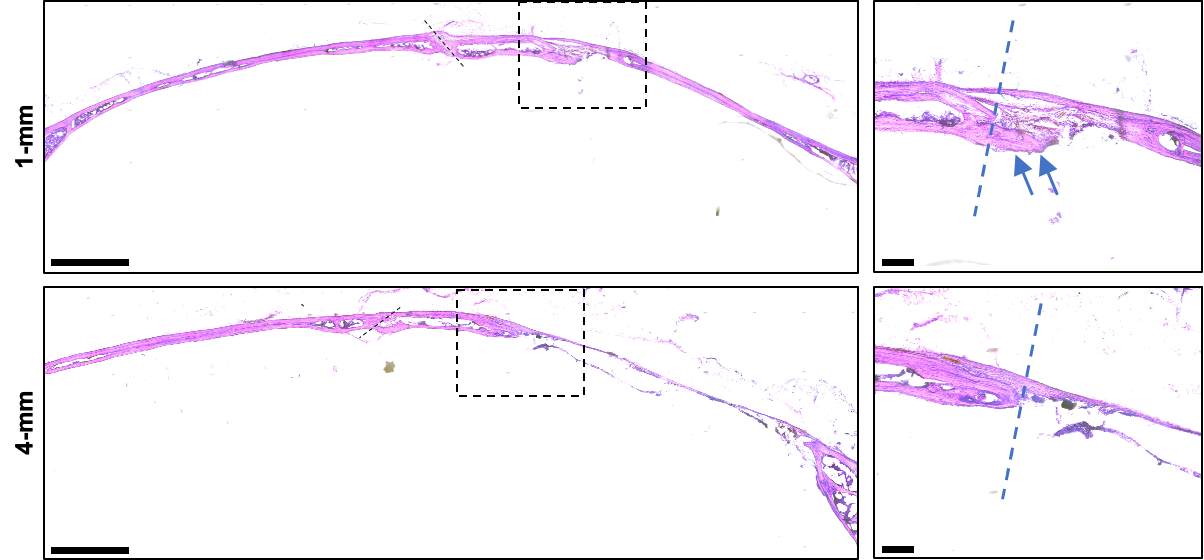

**Supplemental Figure 1: Calvarial cross-sections from sub-critical (1-mm) and critical (4-mm) calvarial defects.** H&E staining of coronal calvarial cross sections at week 8 after injury. Black dashed line represents the sagittal suture region. Dashed boxes demonstrate region of high magnification, which contains the medial osteogenic front for all samples. N=3 mice per group. Bone defect edge is indicated by blue dashed line. Bony ingrowth is labeled by blue arrows. Scale bars: 1000 μm (left) and 100 μm (right).

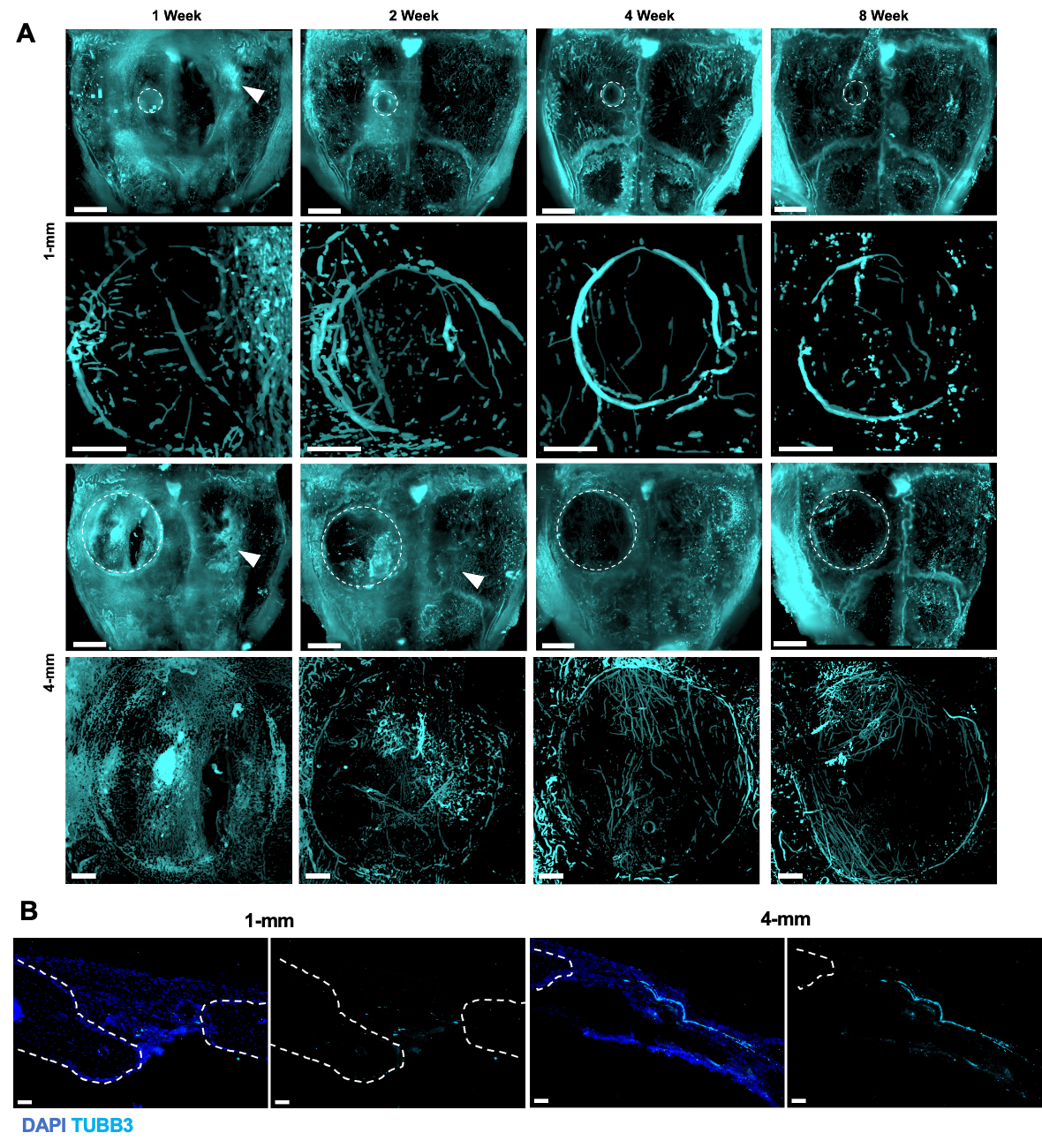

**Supplemental Figure 2: Innervation patterns in healing and non-healing calvarial defects.** A) MIP of TUBB3+ nerves imaged by QLSM at week 1, 2, 4, and 8 after injury. Dashed lines represent defect regions. White arrows demonstrate contralateral response at early timepoints. Overall calvaria scale bar is 100 μm. 1-mm scale bar: 100 μm. 4-mm scale bar: 300 μm. B) MIP of TUBB3 (cyan) and DAPI (blue) 30 μm calvarial cross-section. Region of interest includes the medial osteogenic front of the defect. Dashed lines represent the bone edge. Scale bar: 100 μm.

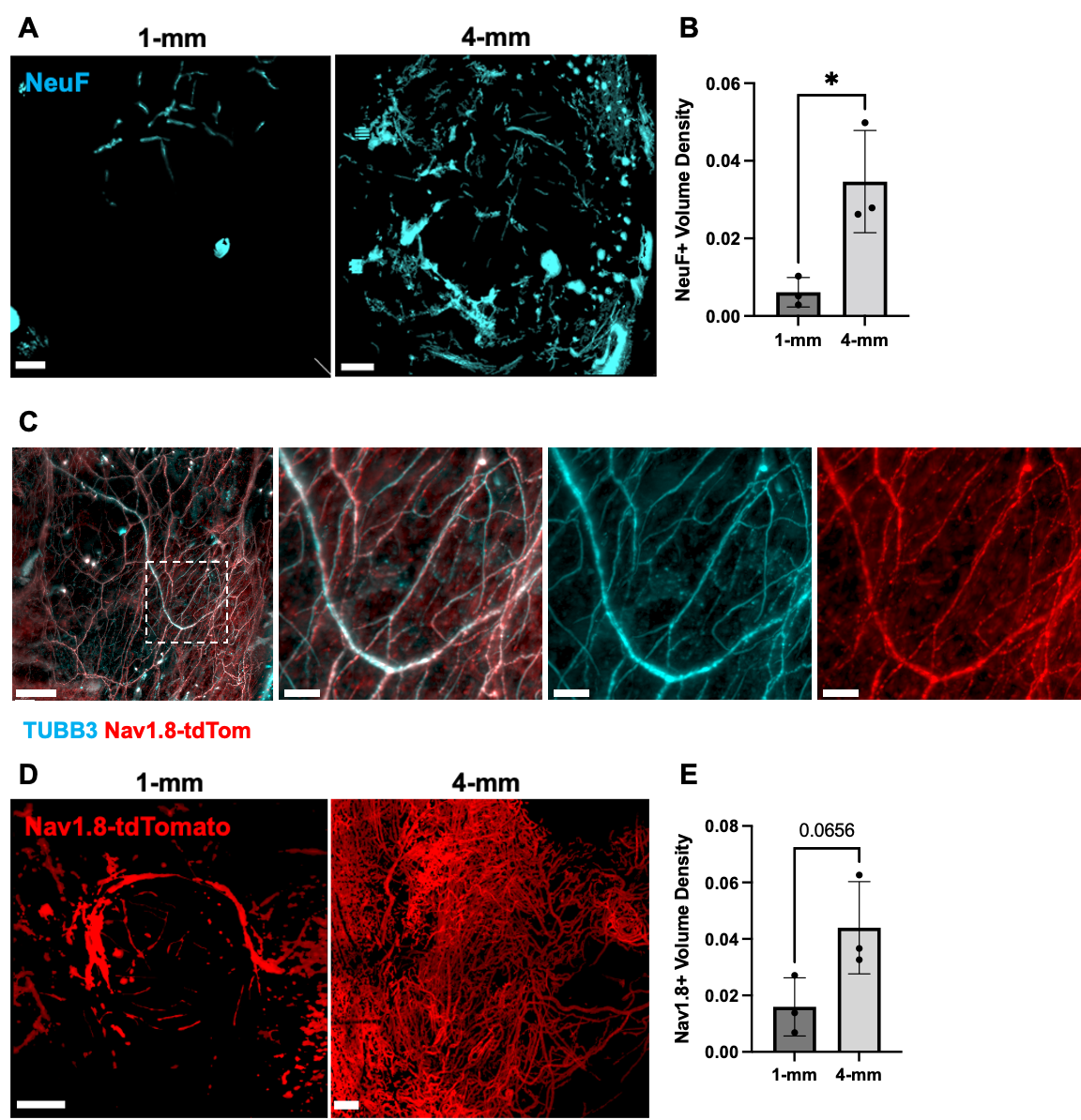

**Supplemental Figure 3: Sensory nerve staining in sub-critical (1-mm) and critical (4-mm) calvarial defects.** A) MIP of NeuF+ nerves imaged by QLSM at week 8 after injury. 1-mm scale bar: 100 μm. 4-mm scale bar: 300 μm. B) NeuF+ volume density quantification. C) A NaV1.8-tdTomato reporter mouse was used in combination with TUBB3 immunohistochemistry (cyan), imaged by QLSM in the uninjured calvaria. Dashed lines present region of high magnification. Scale bar: 400 μm. High magnification region scale bar: 100 μm. D) MIP of NaV1.8-tdTomato nerves imaged by QLSM at week 8 after injury. 1-mm scale bar: 200 μm. 4-mm scale bar: 300 μm. G) NaV1.8+ volume density quantification. Data are Mean ± 1 SD. p* < 0.05. N=3.

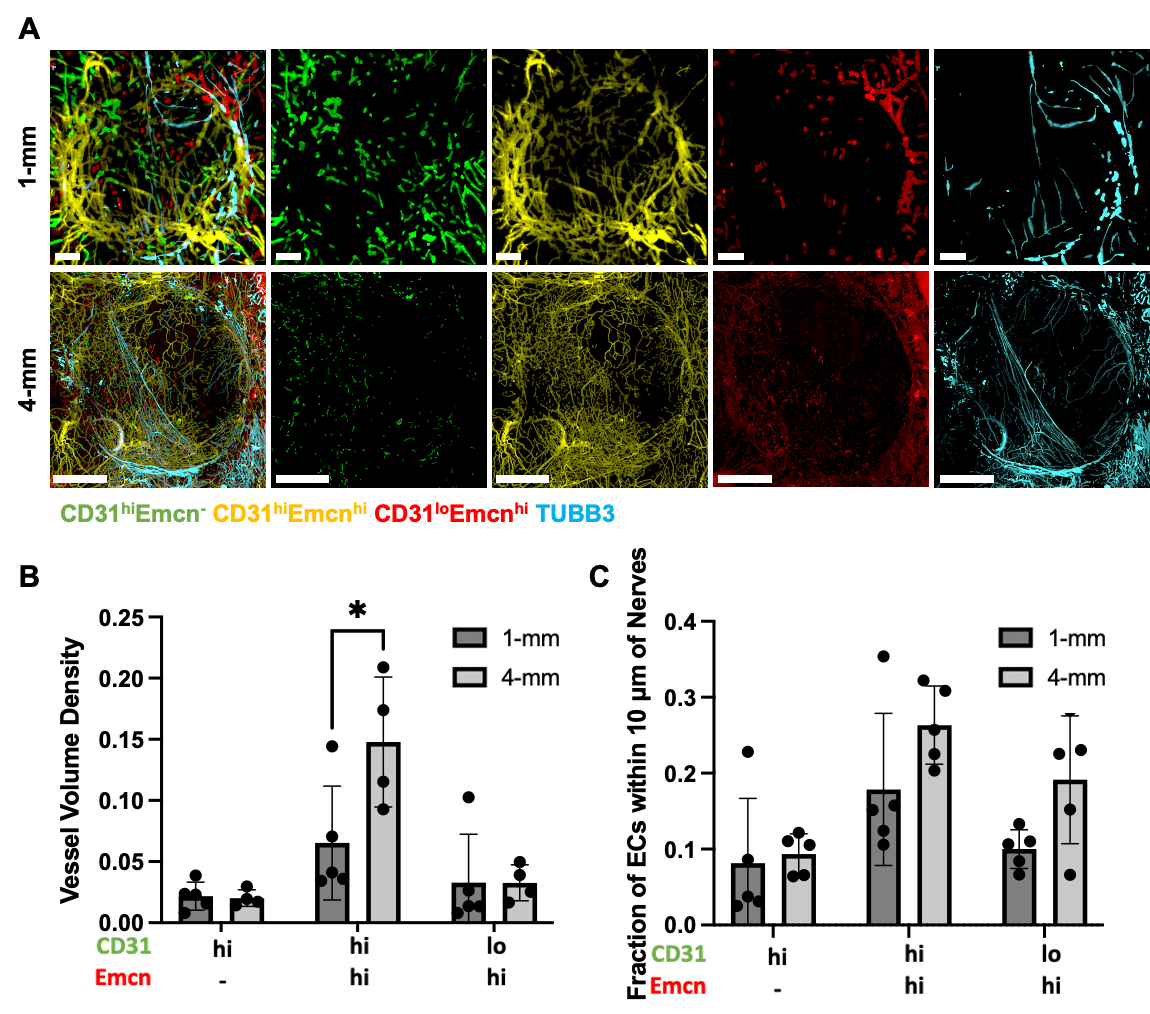

**Supplemental Figure 4: Neurovasculature of healing and non-healing calvarial defects.** A) MIP of TUBB3+ nerves (cyan) and CD31^hi^Emcn^-^ (green), CD31^hi^Emcn^hi^ (yellow), and CD31^lo^Emcn^hi^ (red) blood vessels imaged by QLSM. 1-mm scale bar: 100 μm. 4-mm scale bar: 1000 μm. B) Vessel volume density quantification. C) Quantification of fraction of each endothelial cell (EC) phenotype within 10 μm of nerves. Data are Mean ± 1 SD. p* < 0.05. N=5.

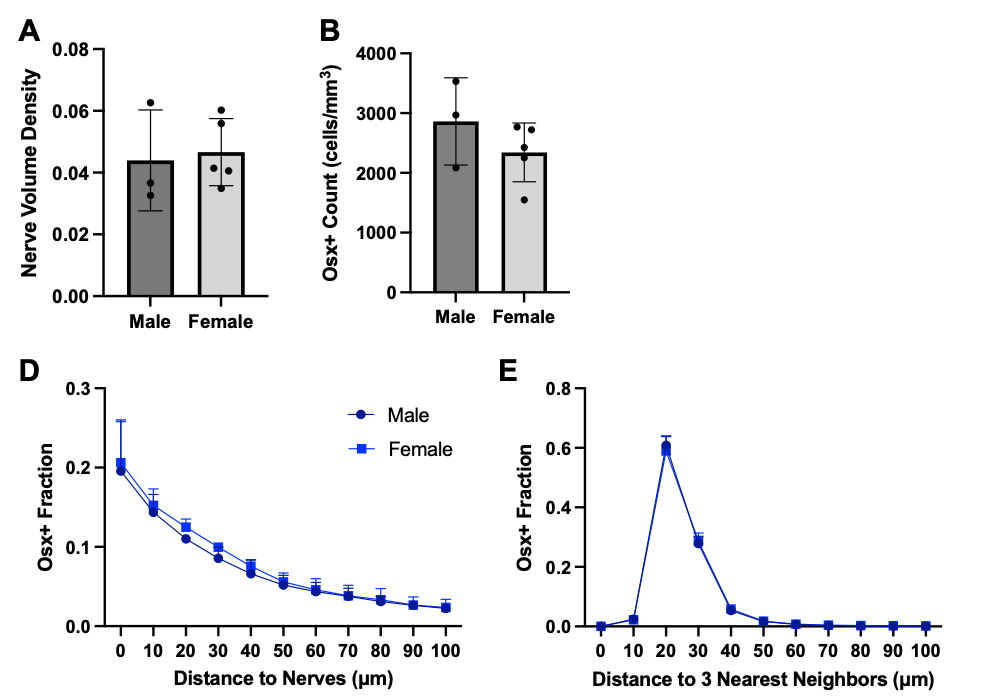

**Supplemental Figure 5: Comparison of nerve volume density and osteoprogenitor fraction after injury in male and female animals.** A) NaV1.8-tdTomato+ volume density quantification. B) Osx+ count quantification. C) Histogram of Osx+ osteoprogenitor spatial association with NaV1.8-tdTomato+ nerves. D) Histogram of Osx+ osteoprogenitor distance to three nearest neighbors. Data are Mean ± 1 SD. N=3-5.

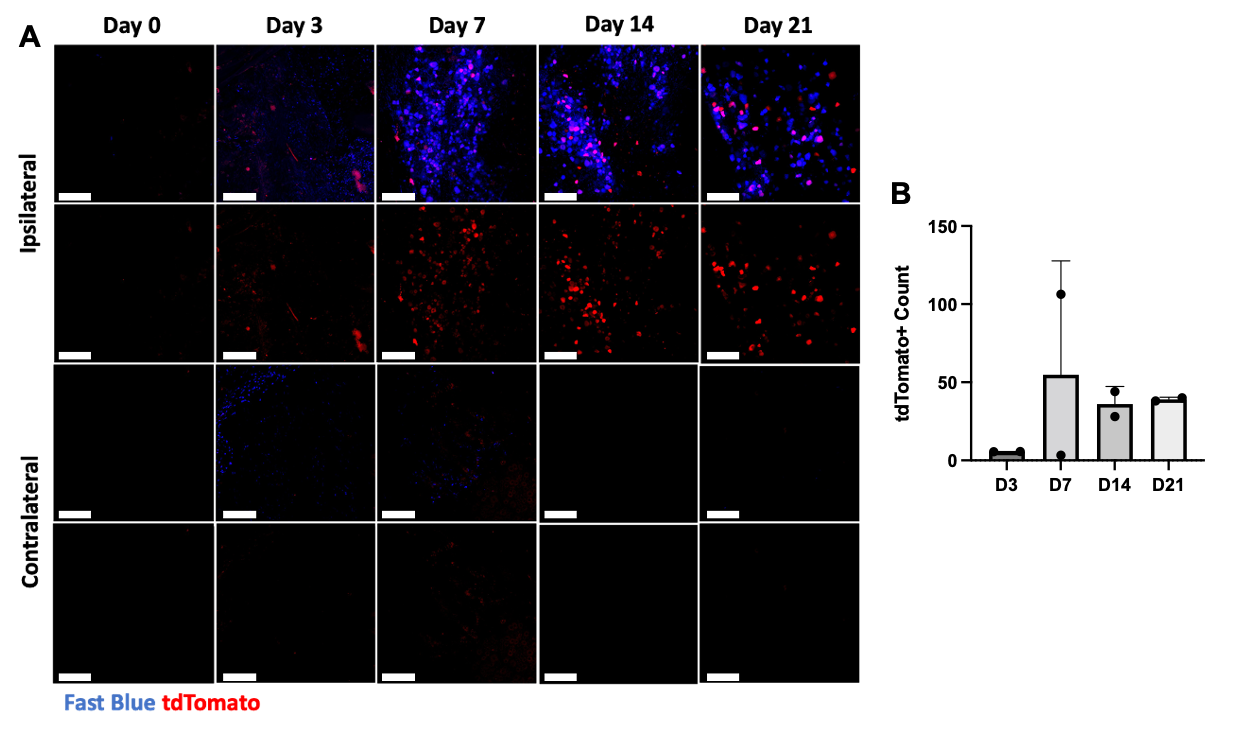

**Supplemental Figure 6: Validation of retrograde tracing.** A) Whole mount imaging of trigeminal ganglia after retrograde tracing using AAV-tdT and fast blue after injection into the mid parietal bone periosteum . B) Quantification of tdTomato+ cell count from whole mount imaging. Data are Mean ± 1 SD. N=2.

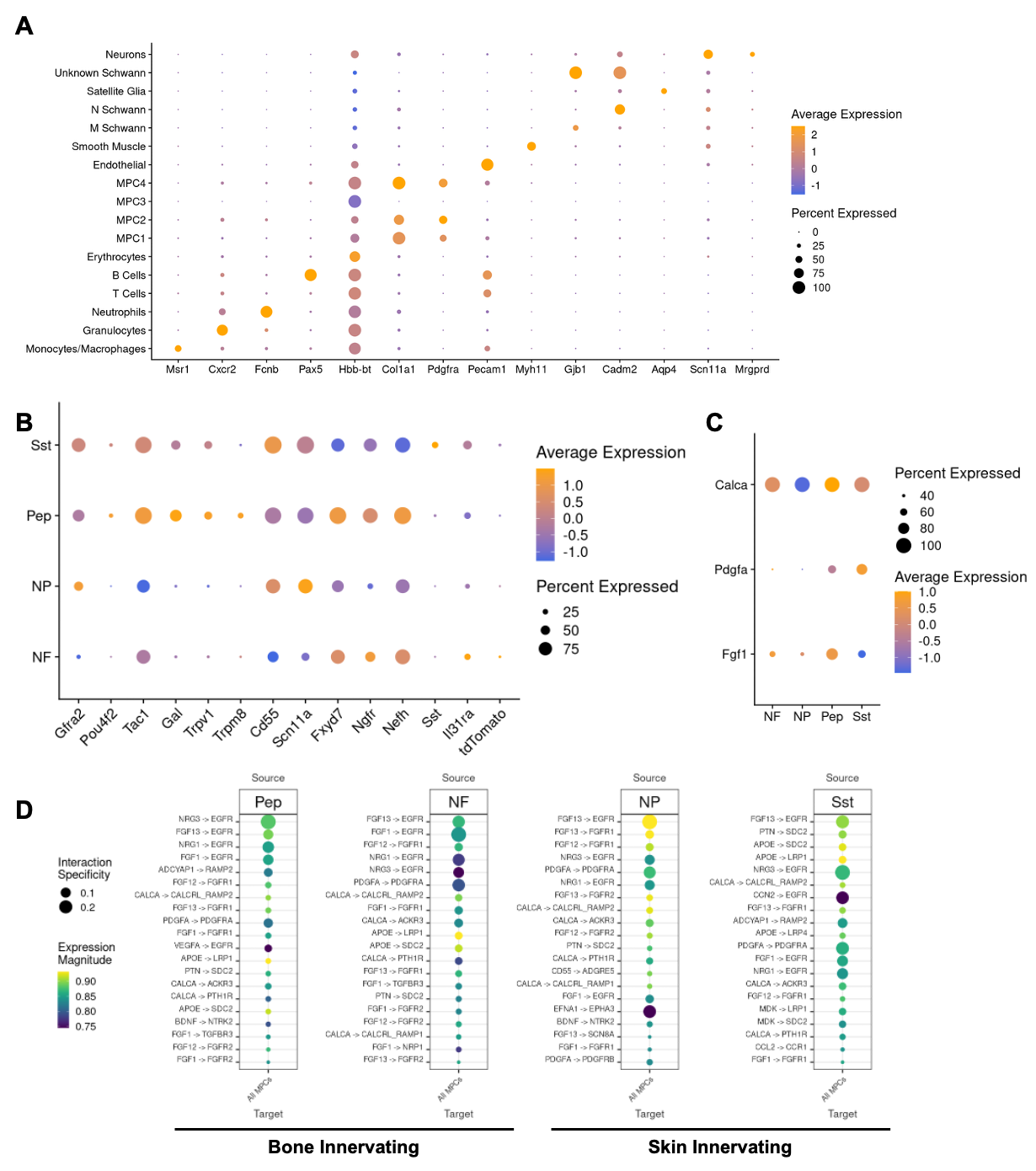

**Supplemental Figure 7: Calvarial nerves express ligands from the FGF, PDGF, and neuropeptide families.** A) DotPlot of cell type specific markers for cluster identification. N Schwann = Non-Myelinating Schwann Cell. M Schwann = Myelinating Schwann Cell. MPC = Mesenchymal Progenitor Cell. B) DotPlot of neuronal specific markers used for cluster identification. C) DotPlot of ligand enrichment across the identified neuronal subpopulations. D) LIANA ligand receptor pair analysis across all neuronal subpopulations to MPCs combined.

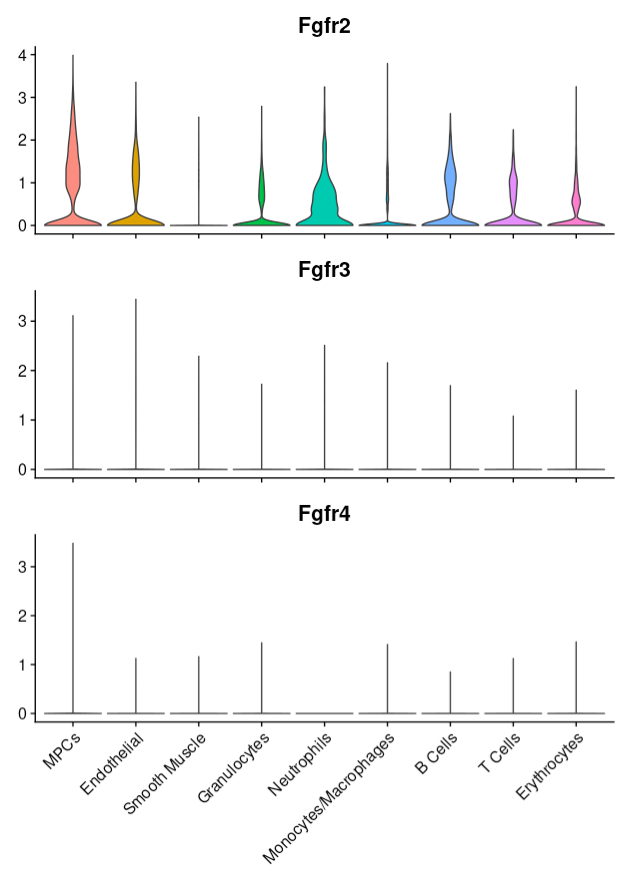

**Supplemental Figure 8: FGFR expression across clusters.** FGFR2. FGFR3, FGFR4 expression across clusters across for all integrated timepoints.

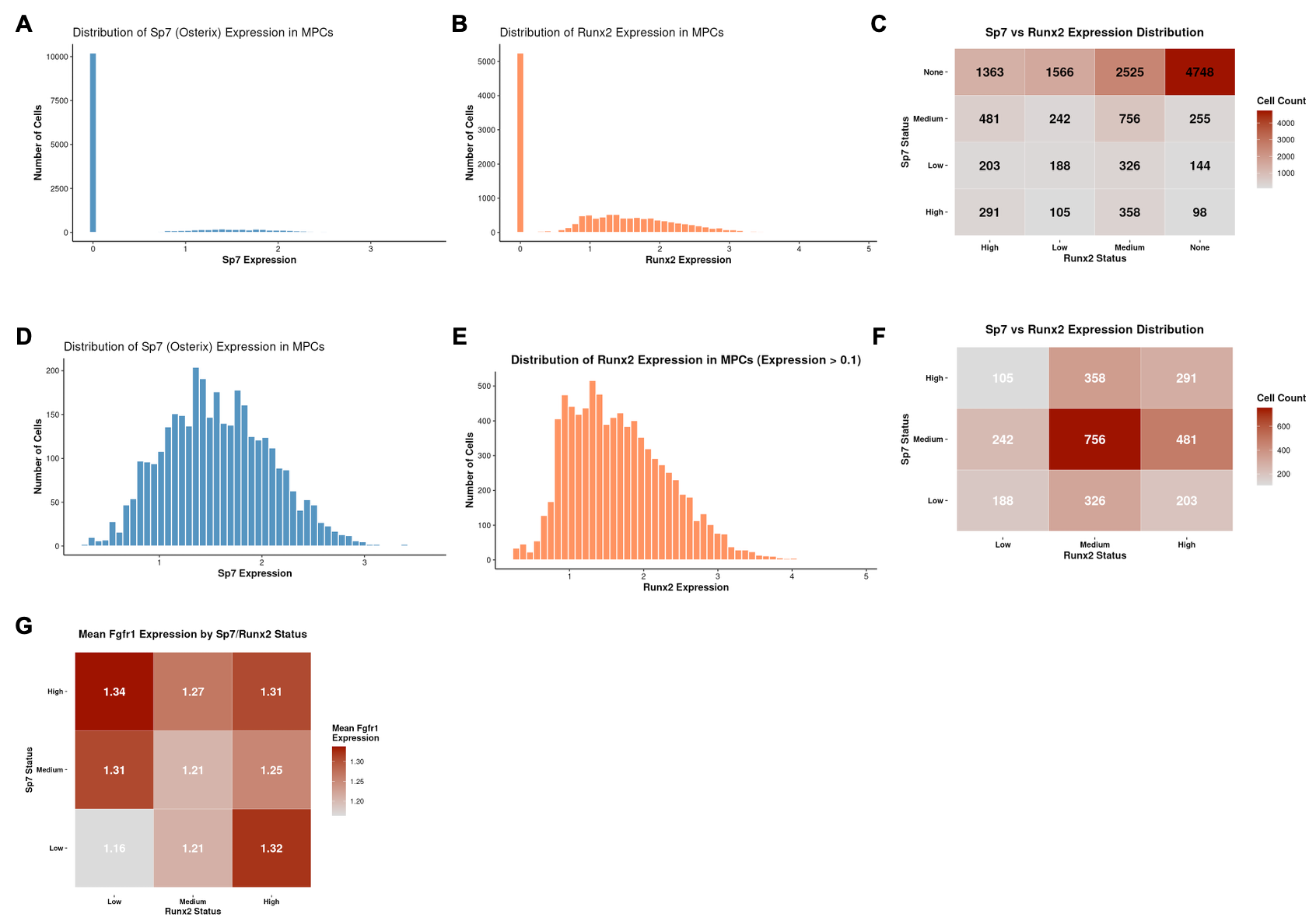

**Supplemental Table 1.** List of all key antibodies used in this study.

**Supplemental Figure 9: FGFR1 expression in Sp7 and Runx2 osteoprogenitors.** A) Histogram distributions of Sp7 (Osterix) expression in full Col1a1-positive MPC population. B) Histogram distributions of Runx2 expression in full Col1a1-positive MPC population. C) Cell distribution heatmap across all Sp7 and Runx2 expression bins, including non-expressing populations. D) Histogram distributions of Sp7 expression in MPCs after filtering out cells with expression ≤0.1. E) Histogram distributions of Runx2 expression in MPCs after filtering out cells with expression ≤0.1. F) Cell distribution heatmap across Sp7 and Runx2 expression bins (Low, Medium, High) in the filtered expressing population. Cells were categorized based on quartile thresholds: Sp7 Low (0.1-1.19), Medium (1.19-1.93), High (>1.93); Runx2 Low (0.1-1.148), Medium (1.148-2.03), High (>2.103). G) Fgfr1 mean expression heatmap across each Sp7/Runx2 expression bin.

| **Antibody** | **Source** | **Identifier** |
| --- | --- | --- |
| ***Immunofluorescence*** | | |
| Rabbit anti-mouse TUBB3 (1:200) | Abcam | ab18207 |
| Rabbit anti-mouse NeuF (1:200) | Thermo Fisher Scientific | PA3-16721 |
| Goat anti-mouse/rat CD31 (1:200) | R&D Systems | AF3628 |
| Rat anti‐mouse/rat Emcn (1:50) | Santa Cruz Biotechnology | sc‐65495 |
| Goat anti-tdTomato (1:200) | SicGen | AB8181 |
| Rabbit anti-mouse Osx (1:200) | Abcam | ab209484 |
| Rabbit anti-mouse OCN (1:200) | Abcam | ab93876 |
| Mouse anti-mouse/human FGF1 (1:100) | Abcam | ab169748 |
| Rabbit anti-mouse FGFR1 (1:200) | Sigma Aldrich | HPA056402 |
| Donkey anti‐goat AF800 plus, 0.67 mg/mL (1:100) | Thermo Fisher Scientific | A32930 |
| Donkey anti‐rabbit AF647 plus, 0.67 mg/mL (1:300) | Thermo Fisher Scientific | A32795 |
| Donkey anti‐goat biotin, 0.75 mg/mL (1:200) | Thermo Fisher Scientific | A16009 |
| Goat anti-mouse Multi-rAb™ CoraLite® Plus 647 | Proteintech | RGAM005 |
| Streptavidin AF555 conjugate, 0.67 mg/mL (1:200) | Thermo Fisher Scientific | S32355 |
